# Memantine Inhibits Calcium-Permeable AMPA Receptors

**DOI:** 10.1101/2024.07.02.601784

**Authors:** Elisa Carrillo, Alejandra Montaño Romero, Cuauhtemoc U. Gonzalez, Andreea L. Turcu, Shao-Rui Chen, Hong Chen, Hui-Lin Pan, Santiago Vázquez, Edward C. Twomey, Vasanthi Jayaraman

**Affiliations:** Center for Membrane Biology, Department of Biochemistry and Molecular Biology, University of Texas Health Science Center at Houston, Houston, TX 77030, USA; MD Anderson Cancer Center UTHealth Graduate School of Biomedical Sciences, University of Texas Health Science Center at Houston, Houston, TX, 77030, USA; Laboratori de Química Farmacèutica (Unitat Associada al CSIC), Facultat de Farmàcia i Ciències de l’Alimentació i Institut de Biomedicina (IBUB), Universitat de Barcelona, Av. Joan XXIII, 27-31, 08028 Barcelona, Spain; MD Anderson Cancer Center, Houston, TX, 77030, USA; Department of Biophysics and Biophysical Chemistry, Johns Hopkins University School of Medicine, Baltimore, MD, 21205, USA; Solomon H. Snyder Department of Neuroscience, Johns Hopkins University School of Medicine, Baltimore, MD, 21205, USA; The Beckman Center for Cryo-EM at Johns Hopkins, Johns Hopkins University School of Medicine, Baltimore, MD, 21205, USA; Diana Helis Henry Medical Research Foundation, New Orleans, LA, 70170, USA

## Abstract

Memantine is an US Food and Drug Administration (FDA) approved drug that selectively inhibits NMDA-subtype ionotropic glutamate receptors (NMDARs) for treatment of dementia and Alzheimer’s. NMDARs enable calcium influx into neurons and are critical for normal brain function. However, increasing evidence shows that calcium influx in neurological diseases is augmented by calcium-permeable AMPA-subtype ionotropic glutamate receptors (AMPARs). Here, we demonstrate that these calcium-permeable AMPARs (CP-AMPARs) are inhibited by memantine. Electrophysiology unveils that memantine inhibition of CP-AMPARs is dependent on their calcium permeability and the presence of their neuronal auxiliary subunit transmembrane AMPAR regulatory proteins (TARPs). Through cryo-electron microscopy we elucidate that memantine blocks CP-AMPAR ion channels in a unique mechanism of action from NMDARs. Furthermore, we demonstrate that memantine reverses a gain of function AMPAR mutation found in a patient with a neurodevelopmental disorder and inhibits CP-AMPARs in nerve injury. Our findings alter the paradigm for the memantine mechanism of action and provide a blueprint for therapeutic approaches targeting CP-AMPARs.

---

Ionotropic glutamate receptors are the primary mediators of excitatory transmission in the mammalian central nervous system (*1, 2*). They are broadly classified into four subtypes: amino-3-hydroxy-5-methyl-4-isoxazolepropionic acid (AMPA), N-methyl-D-aspartate (NMDA), kainate, and delta receptors (*1, 3*). AMPA receptors (AMPARs) mediate fast synaptic signaling and are predominantly calcium impermeable (*1, 2*). AMPARs are formed by combinations of four subunits: GluA1, GluA2, GluA3, and GluA4, which can assemble as homomeric or heteromeric combinations (*1*). GluA2 is unique among these subunits as it exists predominantly in an edited form where the glutamine residue at site 607 (referred to as the Q/R site) is edited to arginine (*4, 5*). The Q/R edited version of GluA2 confers calcium impermeability in AMPARs. Most AMPARs in the mammalian central nervous system contain this edited GluA2 subunit, making them mostly calcium impermeable (*1*).

However, in various neuropathological conditions, there is an increase in the fraction of calcium-permeable AMPARs (CP-AMPARs). Specifically, down-regulation in RNA editing at the Q/R site of GluA2 has been associated with Alzheimer’s disease (*6*), sporadic and familial amyotrophic lateral sclerosis (*7*), seizure vulnerability (*8*), and in malignant gliomas (*9*). Additionally, a decrease in overall GluA2 subunits and an increase in the proportion of CP-AMPARs formed from a combination of GluA1, GluA3, and GluA4 receptors have been shown in gestational hypoxia (*10*), neuropathic pain (*11*), prion protein-mediated excitotoxicity (*12*), an ALS model (*13*), and a mouse model of glaucoma (*14*). The changes in GluA2 editing and/or protein levels lead to an increase of CP-AMPARs in these neurological disorders, highlighting the need for pharmacological agents that can inhibit their activity.

Memantine (Fig. 1A) has been previously thought to be a selective inhibitor of NMDA receptors (NMDARs), with no significant effect on AMPARs (*15, 16*). Memantine is a US-FDA approved drug for treatment of Alzheimer’s and dementia. Memantine inhibits NMDARs by acting as a blocker of the NMDAR ion channel (*15, 16*). Given the increased appreciation for CP-AMPARs in neurological diseases, we hypothesized that memantine may have polypharmacology and also act through inhibiting CP-AMPARs. The lack of memantine inhibition for AMPARs in prior experiments may be because they were studied in a neuronal culture where there are predomintantly calcium-impermeable AMPARs (CI-AMPARs) (*16*). Furthermore, the initial electrophysiology studies on AMPARs in over-expressed systems were performed in the absence of auxiliary subunits, while physiological AMPARs are associated with auxiliary subunits (*17, 18*). These auxiliary subunits alter the biophysical and structural properties of the receptors (*2, 18–24*). In particular, the highly prevalent auxiliary subunits trasmembrane AMPAR regulatory proteins (TARP)-γ2 and -γ8, stabilize the open state of the receptor (*2, 20, 23*).

**Figure 1.**
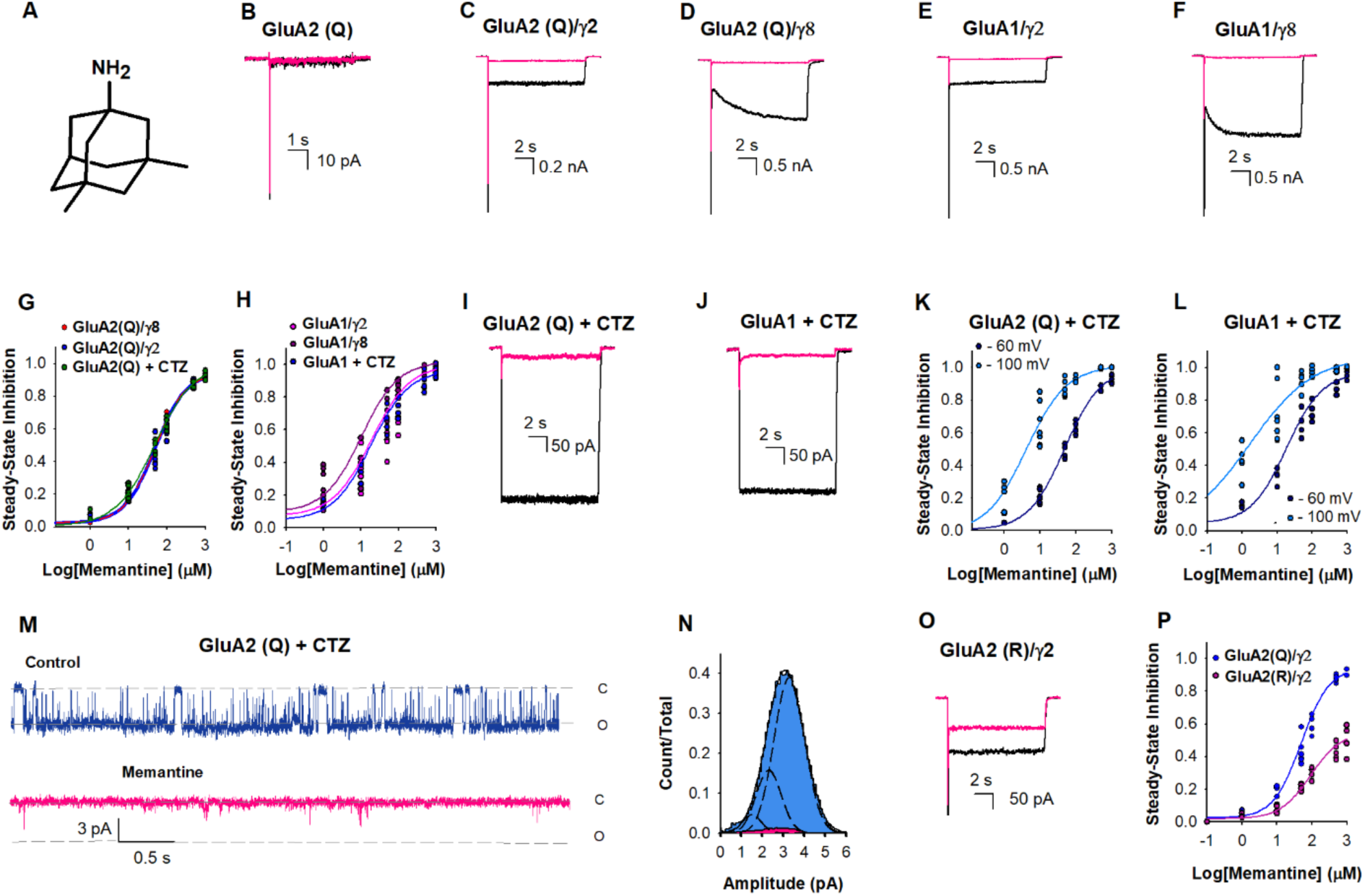
Memantine inhibition of AMPA receptors. (A) Chemical structure of memantine. Representative current traces due to 10 mM glutamate in the absence (black) and presence of 500 µM memantine (pink) from HEK-293 cells expressing (B) GluA2(Q), (C) GluA2(Q)/γ2, (D) GluA2(Q)/γ8, (E) GluA1/γ2, and (F) GluA1/γ8. (G) The dose-dependent inhibitory effects of memantine on GluA2 (Q)/γ2 with IC_50_ 49 ± 2 µM (•), GluA2 (Q)/γ8 with IC_50_ 49 ± 3 µM (•), and GluA2 (Q) in the presence of CTZ with IC_50_ 48 ± 3 µM (•). Each dot represents data from a different cell. (H) The dose-dependent inhibitory effects of memantine on GluA1/γ2 with IC_50_ 15 ± 2 µM (•), GluA1/γ8 with IC_50_ 10 ± 2 µM (•) and GluA1 in the presence of CTZ with IC_50_ 17 ± 3 µM (•). (I) Representative current trace for GluA2(Q) traces due to 10 mM glutamate in the presence of 100 μM CTZ and in the absence (black) and presence of 500 µM memantine, and (J) GluA1 traces due to 10 mM glutamate in the presence of 100 μM CTZ and in the absence (black) and presence of 500 µM memantine. (K) The dose-dependent memantine inhibition on GluA2(Q) in the presence of CTZ at -60 (•) and -100 mV (•), with IC_50_ 48 ± 3 µM and 4.1 ± 1.3 µM, respectively. Each dot represents data from a different cell. (L) The dose-dependent memantine inhibition on GluA1in the presence of CTZ at -60 (•) and -100 mV (•), with IC_50_ 17 ± 3 µM and 2.3 ± 0.3 µM, respectively. Each dot represents data from a different cell. (M) Single-channel currents were recorded from GluA2(Q) in the presence of 100 μM CTZ during continuous application of 10 mM glutamate alone (blue) and in the presence 500 µM of memantine (pink). Openings are shown as downward deflections. (N) Amplitude histogram from the single channel recordings showing conductance (n= 5). (O) Representative current trace for GluA2(R)/γ2 due to 10 mM glutamate in the absence (black) and presence of 500 µM memantine. (P) The dose-dependent inhibitory effects of memantine on GluA2 (R)/γ2 with IC_50_ 1129 ± 85 µM (•), compared to GluA2 (Q)/γ2 with IC_50_ 49 ± 2 µM (•).

Here we show that memantine inhibits CP-AMPARs in micromolar concentrations through electrophysiology and cryo-electron microscopy (cryo-EM). While CP-AMPARs are inhibited at tens of micromolar concentration, CI-AMPARs are inhibited only at hundreds of micromolar concentrations of memantine even in the presence of auxiliary subunits γ2 and γ8. We also show that memantine more effectively inhibits CP-AMPARs containing patient mutation at the ion channel selectivity filter. This mutatation cause significant neurodevelopmental disorders (*25*), and thus memantine has a potential utility in inhibiting the gain-of-function seen for these mutations. Cryo-EM of activated CP-AMPARs in the presence of memantine shows that memantine directly interacts with the AMPAR Q/R site while sitting in the hydrophobic pocket of the ion channel, and inhibits CP-AMPARs through rearrangement of the selectivity filter. This mechanism is unique from polyamine-based pore blockers of CP-AMPARs. Finally, we show that memantine inhibits CP-AMPARs in a nerve injury pain model. Our findings uncover that memantine inhibits CP-AMPARs, how inhibition occurs, show that memantine may be an effective treatment in AMPAR-based disorders, and provide new foundations for therapeutic design.

## Results

### Memantine inhibition CP-AMPARs require auxiliary subunits

We recorded whole-cell currents induced by 10 mM glutamate from HEK-293 cells expressing the CP-AMPARs, homomeric GluA2(Q), and homomeric GluA1, under various conditions in the presence and absence of memantine. When GluA2(Q) receptors were studied in isolation the whole-cell currents induced by 10 mM glutamate did not show any significant inhibition even with 500 µM memantine (Fig. 1B). However, when auxiliary subunits γ2 (Fig. 1C) and γ8 (Fig. 1D) were present, both the peak and steady-state currents of GluA2(Q) receptors were inhibited by 500 µM memantine. Memantine also inhibited CP-AMPAR homomeric GluA1 in the presence of auxiliary subunits γ2 (Fig. 1E) and γ8 (Fig. 1F). The IC_50_ value for memantine inhibition for GluA2(Q) in the presence of γ2 was similar to that in the presence of γ8, 49 ± 2 µM and 48 ± 3, respectively (Fig. 1G). The IC_50_ values for the inhibition of GluA1 were lower than GluA2(Q), being 15 ± 2 µM and 10 ± 2 µM, in the presence of γ2 and γ8, respectively (Fig. 1H).

The presence of auxiliary subunits γ2 or γ8 stabilizes the AMPAR open state and reduces the rate and extent of desensitization seen in GluA2(Q) receptors. Thus the inhibition by memantine under these conditions suggests that memantine inhibits the open channel form of the receptor. To test this idea, we studied inhibition by memantine of GluA2(Q) and GluA1 receptors stabilized in the open channel state using 100 μM cyclothiazide (CTZ), a positive allosteric modulator (Fig. 1I and Fig. IJ). Under these conditions, 500 µM memantine showed inhibition of the steady-state currents, consistent with the findings in experiments with γ2 and γ8, providing further confirmation that memantine can inhibit GluA2(Q) receptors when the open channel state of the receptor is stabilized. For GluA2(Q), the IC_50_ for memantine block in the presence of CTZ was 48 ± 3 µM (Fig. 1K), and for GluA1 the the IC_50_ was 17 ± 3 µM (Fig. 1L). These values are similar to those observed in the presence of γ2 and γ8, thus supporting the open channel block mechanism (Fig. 1G and Fig. 1H). The extent of inhibition by memantine is also voltage-dependent with higher inhibition at more negative voltages (Fig. 1K and Fig. 1L). Dose-response curves show a decrease in IC_50_ at more negative voltages with IC_50_ being 4.1 ± 1.3 µM at - 100mV relative to 48 ± 3 µM at -60 mV for the GluA2(Q) (Fig. 1K) and 2.3 ± 0.3 µM at -100 mV relative to 17 ± 3 µM for GluA1 receptors (Fig. 1L). Given that the resting potential can vary from -60 mV to -85 mV depending on the neuronal subtype and even within parts of the neuron (*26*), micromolar concentrations of memantine is expected to have a significant inhibition at CP-AMPARs.

To investigate why there was higher inhibition of steady-state current versus the peak current, we measured the time for inhibition and recovery for GluA2(Q) currents in the presence of glutamate, CTZ, and 500 µM of memantine (Fig. S1). Under these conditions, memantine blocked the steady-state current with a time-constant of 24 ± 4 ms for the inhibition and a time-constant of 99 ± 12 ms for recovery. The slower time constants for block are consistent with the higher inhibition seen in the steady state condition relative to the peak currents in GluA2(Q) and GluA1 receptor in the presence of γ2 and γ8 (Fig. 1).

To characterize the inhibition mechanism at the single-channel level we performed outside-out single-channel recordings of GluA2(Q)/γ2. Single channels in the presence of CTZ and a saturating concentration of glutamate (10 mM) are predominantly open and populate the higher conductance levels (Fig. 1M, blue). In the presence of memantine, the openings were brief and populated lower conductance states (Fig. 1M, pink). Overall, the receptor does not populate high conductance states in the presence of memantine (Fig. 1N). These measurements further support the inhibition by binding to the open channel form of the receptor.

We also investigated the inhibition by trimethylmemantine (TMM), a memantine derivative with three methyl groups at the amine and a permanent charge on the amine group. TMM (Fig. S2) shows a lower inhibition on GluA2(Q) receptors stabilized in the open state by CTZ (Fig. S2), relative to memantine under the same condition, with TMM having a higher IC_50_ value of 384 ± 8 µM. Additionally, TMM shows less inhibition at saturating concentrations relative to that observed with memantine (Supplementary Figure S2). These results suggest a possible role of steric hindrance due to the bulky trimethyl group at the amine site as the cause for reduction in the inhibition, further consistent with memantine being in the pore where it is expected to have steric constraints. This is also seen in the classical NMDA inhibition by N-alkyl derivatives of memantine compared to memantine (*27*).

Furthermore, we confirmed that memantine does not significantly inhibit CI-AMPARs. We studied memantine block with the representative CI-AMPAR GluA2(R) in the presence of γ2. These studies show that memantine has only a minor inhibitory effect at 500 µM concentrations (Fig. 1O). Dose-response curves for steady-state inhibition with varying memantine concentrations confirmed these results, indicating that the IC_50_ value for memantine inhibition was 20 times higher for the GluA2(R)/γ2 receptor compared to the GluA2(Q)/γ2 receptor (Fig. 1P). Even at saturating concentrations of memantine, GluA2(R)/γ2 receptors displayed only partial inhibition of the currents mediated by 10 mM glutamate (Fig. 1P).

### Memantine inhibition at CP-AMPARs is unique from NMDARs

To determine if other NMDAR channel blockers inhibited CP-AMPARs, we tested the effect of MK801 and ketamine, both of which are high affinity (nanomolar) channel blocker of NMDARs. We show that even at hundreds of micromolar concentrations MK801 and ketamine have a minimal inhibitory effect on CP-AMPARs (Fig. S3). Thus, the observation of NMDAR blocker polypharmacology with CP-AMPARs may be unique to memantine.

Additionally, memantine has been shown to have two pathways and sites for inhibition in NMDARs, one pathway through an open channel block and a second pathway through the membrane (*27*). To investigate if a similar mechanism occurs in memantine inhibition in CP-AMPARs, we preincubated cells with memantine before activation with 10 mM glutamate in the presence of CTZ and compared the currents at pH 7.4 and pH 9. No significant differences were observed between pH 7.4 and pH 9 (Fig. S4). This suggests that, unlike what is observed in NMDARs, memantine does not block CP-AMPARs through a membrane pathway.

### Mechanism of memantine channel block in CP-AMPARs

To elucidate the memantine channel block mechanism in CP-AMPARs, we used cryo-electron microscopy (cryo-EM) to capture activated CP-AMPARs in the presence of memantine. We used a well-established CP-AMPAR cryo-EM construct (GluA2-γ2_EM_, Methods), which is a covalent fusion construct between GluA2(Q) and γ2. This construct has been extensively validated previously, both functionally and structurally, and has been utilized to study the structural basis of CP-AMPAR channel block (*28–33*).

We prepared samples for cryo-EM by activating GluA2-γ2_EM_ in the presence of glutamate, CTZ, and memantine, which captured both the open channel memantine-blocked state (GluA2-γ2_mem_) and open channel state without memantine (GluA2-γ2_open_) (Methods, Fig. S5, Table 1). GluA2-γ2_mem_ and GluA2-γ2_open_ are largely similar (root mean squared deviation, RMSD = 0.79 Å), with the exception of the transmembrane domain (TMD), where memantine binds. The overall architecture of GluA2-γ2_mem_ is reminiscent of previously-solved GluA2-γ2 structures and native AMPAR complexes: at the core of GluA2-γ2_mem_ are the CP-AMPAR GluA2(Q) subunits arranged in a tetramer, in complex with four γ2 subunits (Fig. 2A). There is an overall “Y” shape of the AMPAR, with the amino-terminal domain (ATD) and ligand binding domain (LBD) comprising the extracellular domain (ECD; Fig. 2A). The ECD in both GluA2-γ2_mem_ and GluA2-γ2_open_ is two-fold symmetric. Below the ECD is the TMD. In GluA2-γ2_open_, the TMD is two-fold symmetric, as expected, similar to the originally solved open state of GluA2-γ2_EM_ (*32*). The GluA2-γ2_mem_ TMD is asymmetric due to a single copy of memantine in the AMPAR TMD (Fig. 2A).

**Figure 2.**
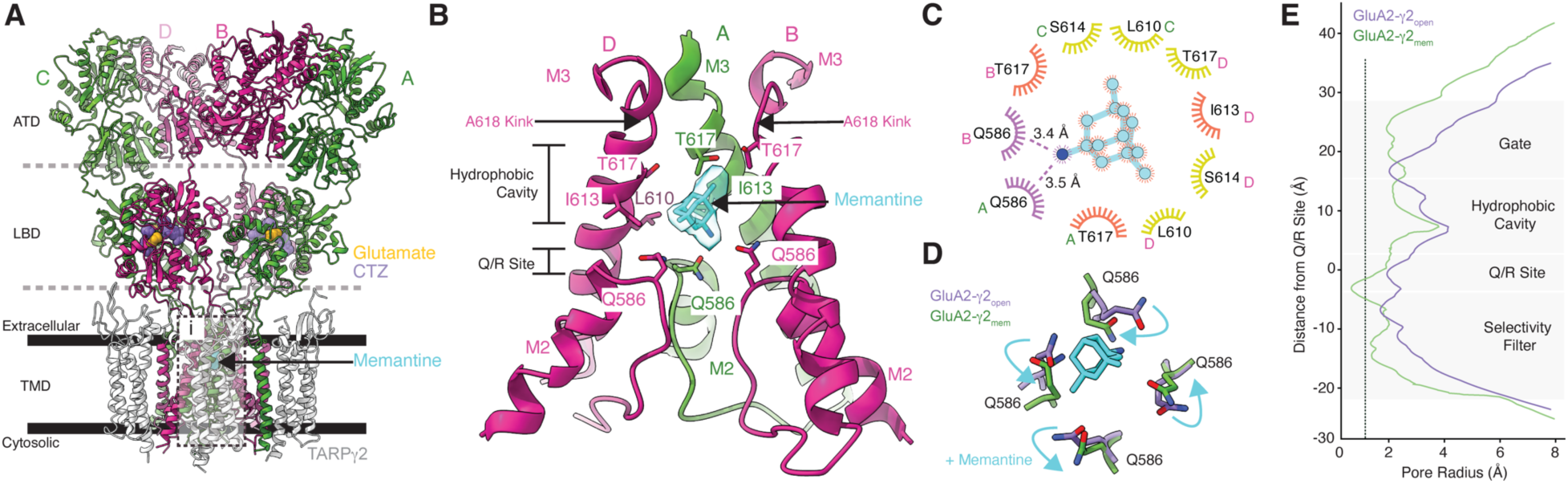
Memantine block revealed by Cryo-EM. (A) Overall architecture of GluA2-γ2_mem_ in a ribbon diagram. GluA2 subunits are labeled depending on their positions (A,C are green; B/D are pink). γ2 subunits are colored white. Dashed lines indiciate domain boundaries, solid bars indicate membrane boundaries. Glutamate (yellow), CTZ (purple), and memantine (cyan) are shown as space filling models. Inset i is where panel B highlights. (B) The GluA2-γ2_mem_ ion channel with M2 and M3 helices. Subunit C is omitted for clarity. The cryo-EM map at the memantine binding site is shown transparently in cyan around the molecule. Crtical binding residues are labeled. Carbon atoms are colored with the color of their respective subunit with oxygen in red, nitrogen in blue. (C) LigPlot of the memantine binding site. Potential polar contacts are indicated with purple eyelashes. Orange eyelashes indicate a distance within 4 Å from memantine. Yellow eyelashes indicate a distance from 4 to 6 Å. (D) How memantine rearranges the Q/R site; cyan arrows indicate changes from GluA2-γ2_open_ (purple) to GluA2-γ2_mem_ (green). (E) Pore radius plots of GluA2-γ2_open_ (purple) and GluA2-γ2_mem_ (green). Dashed line represents the radius of a calcium cation.

The AMPAR ion channel is comprised of the M3 TMD helices and M2 helix, between which is the reentrant loop that contains the Q/R site and selecitivty Filter (Fig. 2B). Memantine binds directly in the CP-AMPAR TMD immediately above the Q/R site, which is the primary determinant of ion channel selectivity in AMPARs. (Fig. 2B). The tricyclodecane backbone of memantine sits in the hydrophobic cavity of the ion channel, and the amine on memantine is directly coordinated by the glutamine residues at the Q/R site (Fig. 2B).

The cryo-EM map of GluA2-γ2_mem_ directly shows the shape of memantine in the ion channel (Fig. 2B). This suggests a singular pose of memantine in the channel, similar to the structure of memantine bound to NMDARs. The memantine binding site is markedly absent from the GluA2-γ2_open_ map (Fig. S6). The memantine binding site in GluA2-γ2_mem_ is resolved to approximately 3-3.5 Å (Fig. S6). Hydrophobic residues in each subunit coordinate the memantine tricyclodecane cage (Fig. 2C). Residues T617 in subunits A and B, as well as I613 in subunit C play the principal roles in coordination via the hydrophobic cavity and are within 4 Å of interaction with memantine. Interestingly, T617 residues are required for calcium coordination in the ion channel (*34*). The presence of memantine at this position likely prevents that possibility. The amine group on memantine is directly coordinated by polar interactions with Q586, the Q607-equivalent in GluA2-γ2_EM_, from subunits A and B. This is reflected in a rearrangement of the Q/R site in GluA2-γ2_mem_ compared to GluA2-γ2_open_ (Fig. 2D). Q586 from the A and B subunit positions move toward the pore center to coordinate memantine, while Q586 at the C and D positions move away from the pore. This overall rearrangement of the Q/R site constricts the remaining selecitivty filter below the Q/R site compared to GluA2-γ2_open_ (Fig. 2E).

The memantine binding site in GluA2-γ2_mem_ directly shows why memantine has increased affinity for CP-AMPARs as opposed to CI-AMPARs. CI-AMPARs contain a bulky, positive charge via editing to ariginine at the Q/R site, which clashes directly with the charged amine group on memantine. And, glutamine at the Q/R site directly coordinates memantine in the pore.

The memantine block mechanism in CP-AMPARs is distinct from polyamine-based blockers. Polyamine blockers such as N,N,N-tri-methyl-5-[(tricyclo[3.3.1.13,7]dec-1-ylmethyl)amino]-1-pentanaminium bromide hydrobromide (IEM-1460), 1-naphthyl acetyl spermine (NASPM), and Argiotoxin-636 (AgTx-636) block CP-AMPAR ion channels through permeating the selectivity filter with their polyamines, and plugging the channel via bulky hydrophobic headgroups that sit in the hydrophobic cavity (Fig. S7). Memantine blocks CP-AMPAR channels by sitting in the hydrophobic cavity, which ablates cation coordination in the upper vestibule of the channel, and by directly interacting with the Q/R site, which narrows the selectivity filter below.

Memantine binds to a similar site in NMDARs but is more closely coordinated by hydrophobic residues in the NMDAR ion channel (*35*). This may account for the discrepancy in affinity between memantine inhibition in NMDARs versus CP-AMPARs. However, in contrast to inhibition in NMDARs, memantine rearranges the Q/R sites in CP-AMPARs, which occludes the channel.

### A CP-AMPAR neurodevelopmental mutation increases the efficacy of memantine

In our cryo-EM data, we observe that memantine directly interacts with the GluA2 Q/R site. We hypothesized that by placing a negative charge at this site, we could dramatically increase the efficancy of memantine inhibition in AMPARs. In fact, there is a *de novo* mutation in a patient with severe developmental delays and Rett-like syndrome at the GluA2 Q/R site where glutamine is mutated to glutamate (GluA2(E); Q607E) (*25*). Because memantine directly interacts with glutamine at the Q/R site, we hypothesized that memantine may have increased inhibition efficacy in GluA2(E). To test this idea, we investigated memantine inhibition of GluA2(E) AMPARs.

The extent of current inhibition for GluA2(E) in the presence of γ2 was notably higher than that observed for GluA2(Q) and GluA2(R) in the presence of γ2 (Fig. 3A). The IC_50_ for memantine inhibition was determined to be 25 ± 2 μM for GluA2(E) in the presence of γ2 (Fig. 3B), significantly lower than the values of 1129 ± 85 μM for the GluA2(R) in the presence of γ2 and 49.3 ± 2.4 μM for GluA2(Q) in the presence of γ2 (Fig. 1I). This is also reflected in the voltage dependence which show that the IC_50_ for memantine GluA2(E) is shifted from 25 ± 2 μM to 5.7 ± 1 μM when the voltage was decreased from -60 mV to -100 mV (Fig. 3B). The observed trends of greater inhibition current extent and a lower IC_50_ value for GluA2(E) compared to GluA2(Q), followed by GluA2(R), indicate that the charge at site 607 influences the extent of memantine inhibition, suggesting that positively charged amine group may reside close to 607 site. We performed outside-out single-channel recordings GluA2(E)/ γ2, to further characterize the inhibition mechanism at the single-channel level. GluA2(E)/ γ2 single channel in the presence of CTZ and a saturating concentration of glutamate (10 mM) predominantly populates the higher conductance levels (Fig. 3C, blue) similar to GluA2(Q) in presence of CTZ (Fig. 1M). In the presence of memantine, the openings were brief and populated lower conductance states (Fig. 3C, pink). Amplitude histograms show that the receptor populates primarily low conductance states in the presence of memantine (Fig. 3D). These measurements support the inhibition by binding to the open channel form of GluA2(E)/ γ2 the receptor similar to that seen for GluA2(Q).

**Figure 3.**
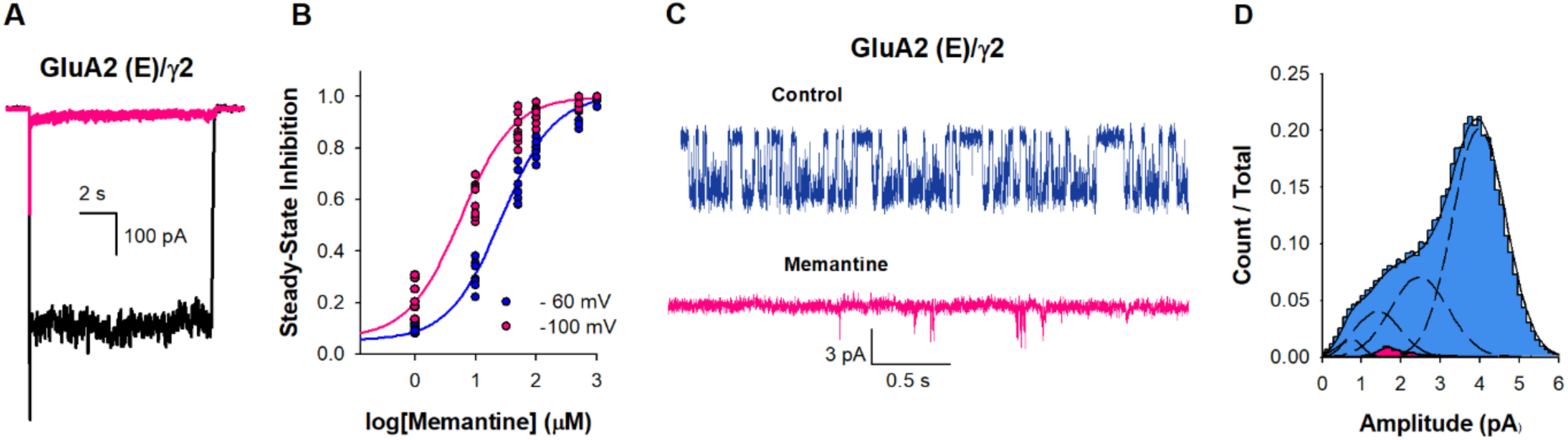
Memantine inhibition of GluA2 (E) /γ2. (A) Representative whole-cell recordings in response to 10 mM glutamate alone (black) or in the presence of 500µM of memantine (pink). (B) The dose dependence of memantine inhibition on GluA2 (E) /γ2 at -60 (•) and -100 (•) mV, with IC_50_ 25 ± 2 µM and 5.7 ± 1 µM, respectively. Each dot represents data from a different cell. (C) Single-channel currents recorded from GluA2 (E) /γ2 during continuous application of 10 mM glutamate alone (blue) and in the presence 500 µM of memantine (pink). Openings are shown as downward deflections. (D) Amplitude histogram from the single channel recordings showing conductance (n = 5).

This data not only supports the mechanism of action observed from our cryo-EM data and functional studies, but also points to the possible utility of memantine in treating AMPAR gain of function in the Q607E mutant.

### Memantine inhibits mEPSCs in cultured neurons

To determine if memantine affected synaptic AMPA receptor signaling, we conducted whole-cell voltage-clamp recordings using high-cell density cortical neuronal cultures. To study mEPSCs, we employed TTX, bicuculline, DL-APV, DCKA and Mg^2+^ in the recording conditions. This ensured that the signaling was mediated by AMPA receptors and NMDA receptors were blocked. In the presence of 10 µM memantine there were no significant changes in the amplitude of spontaneous mEPSCs as well as in the frequency of mEPSCs (Fig. 4). Since the electrophysiological experiments showed inhibition of CI-AMPARs at 500 µM memantine we studied the effect of memantine on mEPSCs at this concentration and showed that memantine does have an effect with a reduction in the amplitude of spontaneous mEPSCs from 28.8 ± 4.6 to 15.4 ± 1.7 pA and a reduction in the frequency of mEPSCs from 21.1 ± 3.1 to 12.3 ± 3.1 (Fig. S8). Pairwise recordings from the same neurons showed that these differences were significant (p= 0.006). These studies show that memantine would not have any effect at tens of micromolar concentration of memantine on the predominant form of AMPA receptor and requires very high concentrations to have any effect.

**Figure 4.**
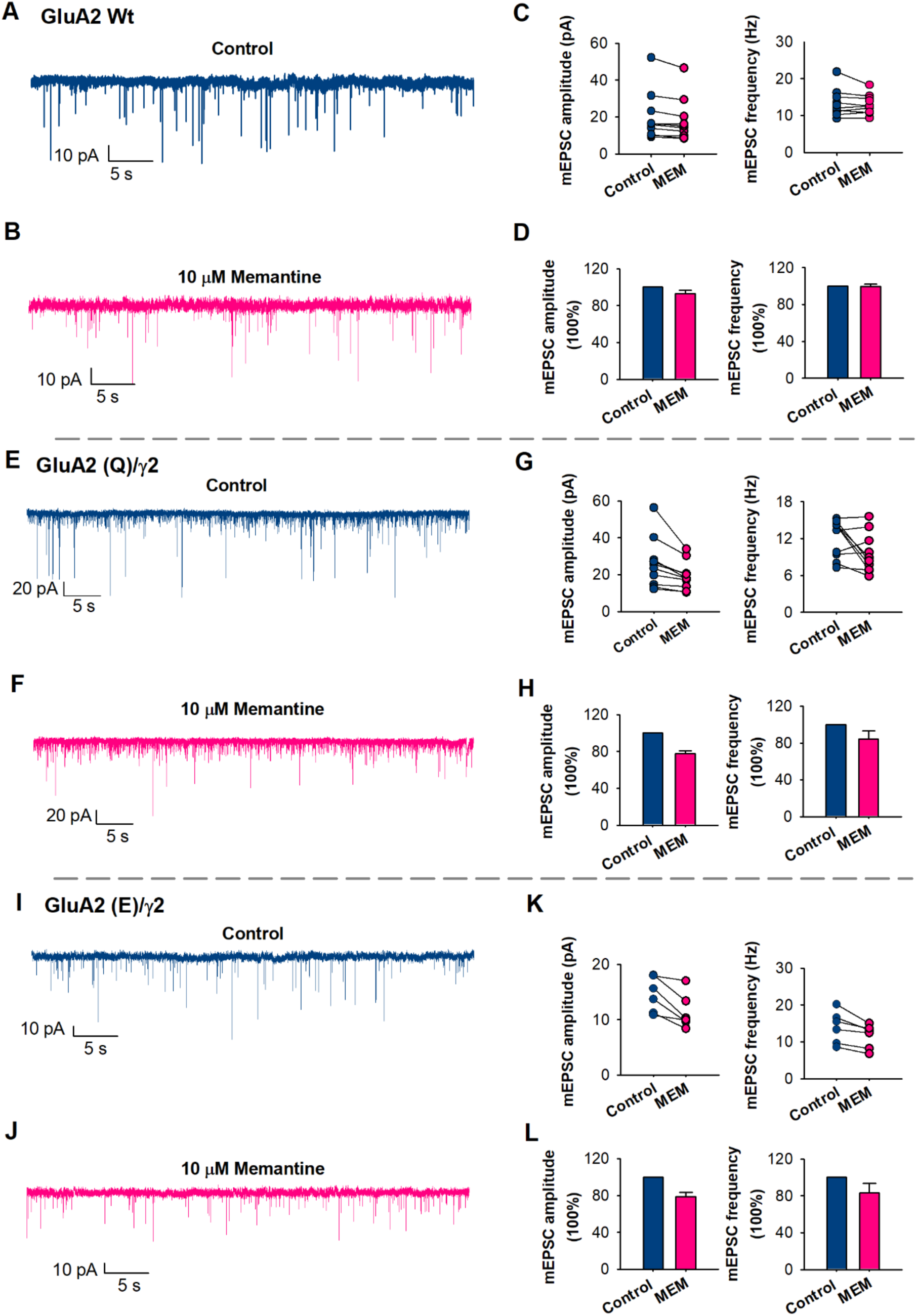
mEPSCs inhibition by Memantine (MEM). (A) Representative spontaneous mEPSCs in native neurons, in control (blue) and (B) in the presence of 10 µM of memantine (pink) (p = 0.7). (C) mEPSC amplitude and frequency were measured from individual neurons. Paired data from each experiment are connected by a line. (D) Bar graphs of the average values of the normalized mEPSC amplitude and frequency, in control (blue) and in the presence of 10 µM of memantine (pink) (n = 10). (E) Representative spontaneous mEPSCs in GluA2 (Q)/γ2 neurons, in control (blue) and (J) in the presence of 10 µM of memantine (pink) (p = 0.02). (G) mEPSC amplitude and frequency from individual neurons. (H) Bar graphs of the average values of amplitude and frequency (n = 10). (I) Representative spontaneous mEPSCs in GluA2 (E)/γ2 neurons, in control (blue) and (J) in the presence of 10 µM of memantine (pink) (p = 0.01). (K) mEPSC amplitude and frequency from individual neurons. (L) Bar graphs of the average values of amplitude and frequency (n = 66).

To determine if memantine would inhibit neurons expressing CP-AMPARs, as is seen in certain pathological states, we transfected wild-type neurons with GluA2(Q)/γ2. Current-voltage curves were used to establish that the neurons are expressing GluA2(Q)/γ2 (Fig. S9). In these neurons we found that10 µM of memantine decreased the amplitude of spontaneous mEPSCs from 25.9 ± 4.8 to 19 ± 2.7 pA and reduced the frequency of mEPSCs from 12.6 ± 1.7 to 10.5 ± 1.1 (Fig. 4). A similar reduction was also observed in neurons transfected with the mutant GluA2(E)/γ2, where 10 µM of memantine decreased the amplitude of spontaneous mEPSCs from 14.6 ± 1.3 to 11 ± 1.2 pA and reduced the frequency of mEPSCs from 14 ± 1.8 to 11.5 ± 1.3 (Fig. 4). Current-voltage curves where used to establish that the neurons are expressing GluA2(E)/γ2 (Fig. S9). These studies show that memantine inhibits CP-AMPARs and the GluA2(E) mutant receptor expressed in neurons at tens of micromolar concentrations similar to what is seen in the HEK-293 cells.

### Memantine inhibits nerve injury-induced synaptic CP-AMPARs

Peripheral nerve injury or painful diabetic neuropathy increases the prevalence of synaptic CP-AMPARs in the spinal dorsal horn, and blocking spinal CP-AMPARs attenuates chronic neuropathic pain (*36, 37*). Although memantine, administered systemically or intrathecally, effectively reduces neuropathic pain in animal models (*38–40*), it has been assumed that its therapeutic action is due to NMDAR blockade. We have demonstrated that α2δ-1 is essential for nerve injury-induced increases in CP-AMPARs (*11*). Given that α2δ-1 exhibits specific expression in VGluT2-expressing excitatory neurons within the spinal cord (*41*), we determined the potential effect of memantine on synaptic CP-AMPARs in genetically labeled spinal VGluT2 neurons of mice subjected to spared nerve injury.

We employed whole-cell voltage-clamp mode to record AMPAR-mediated EPSCs in tdTomato-tagged VGluT2 neurons located in lamina II, evoked monosynaptically from the dorsal root, in the presence of 50 µM AP5, a specific NMDAR antagonist. IEM-1460, recognized as a specific open-channel blocker of CP-AMPARs (*30*), was incorporated at a concentration of 10 mM in the intracellular recording solution to inhibit postsynaptic CP-AMPARs. Before the bath application of memantine, the intracellular analysis of IEM-1460 was allowed for 15 min, which eliminates CP-AMPARs in spinal dorsal horn neurons (*11*). To determine the effect of memantine on CP-AMPARs in the spinal dorsal horn of nerve injured mice, we bath applied memantine at concentrations of 10, 20, and 50 µM for 3 min each, following an ascending order. In VGluT2 neurons recorded with intracellular IEM-1460, memantine at these concentrations failed to produce a significant effect on the amplitude of AMPAR-EPSCs (n = 17 neuron; Fig. 5).

**Figure 5.**
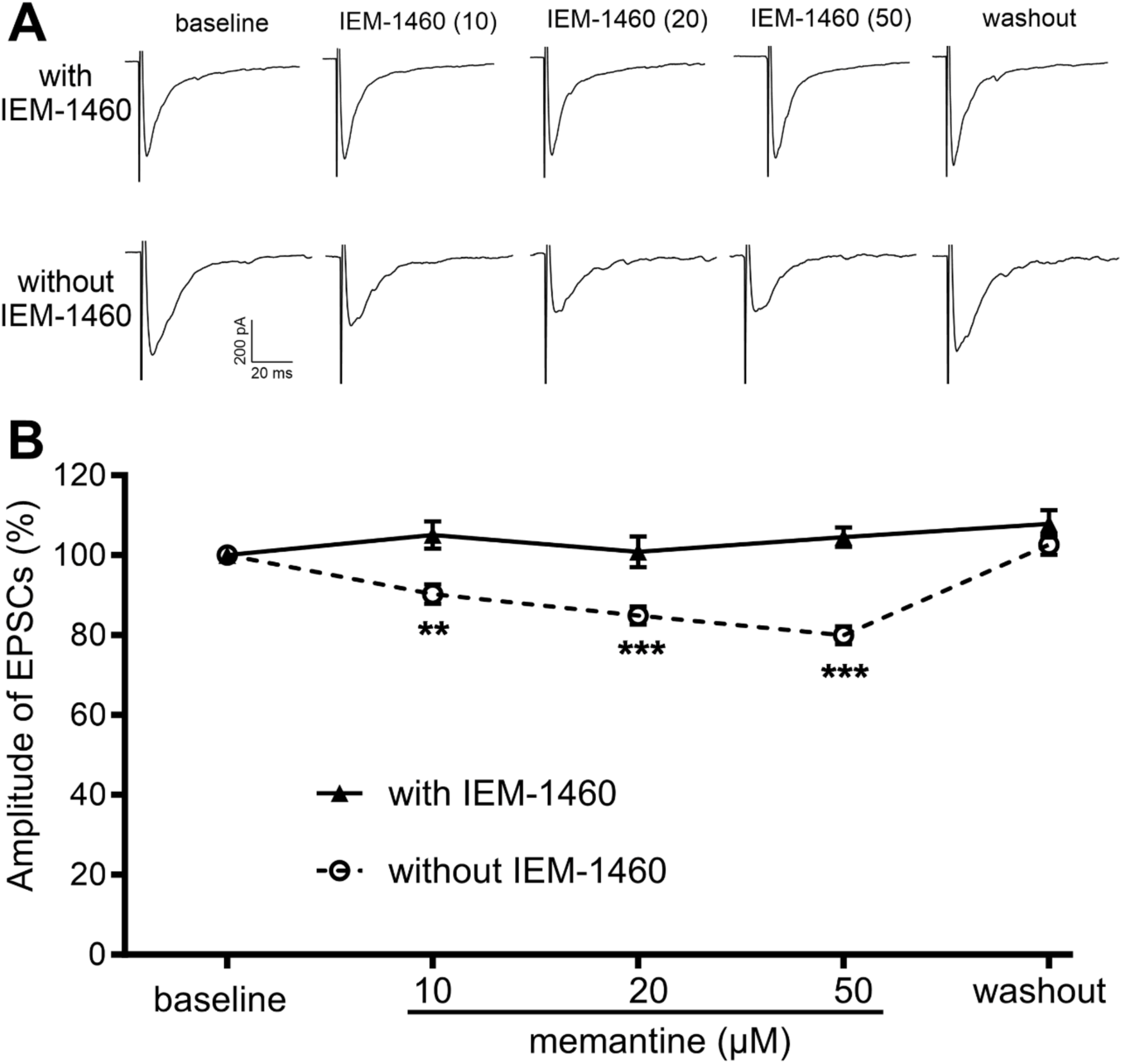
Memantine inhibits synaptic CP-AMPARs in spinal excitatory neurons caused by nerve injury. A and B, Representative recording traces (A) and quantification (B) illustrate the differential effect of bath application of memantine (10, 20, and 50 µM) on monosynaptically evoked AMPAR-EPSCs in spinal VGluT2 neurons of SNI mice recorded with IEM-1460 (n = 17 neurons from 8 mice) and without IEM-1460 (n = 16 neurons from 8 mice). The data were normalized to the baseline value (100%) immediately prior to memantine application. Data are presented as mean ± SEM. **P < 0.01, ***P < 0.001, vs. the baseline control within the group (repeated measures ANOVA followed by Dunnett’s *post hoc* test).

In contrast, in tdTomato-tagged VGluT2 neurons recorded without IEM-1460 in the intracellular solution, bath application of 10, 20, and 50 µM memantine significantly reduced the amplitude of evoked AMPA receptor -EPSCs in a concentration-dependent manner (n = 16 neuron; Fig. 5). The amplitude of evoked AMPAR-EPSCs fully returned to the baseline level ∼10 min after memantine washout. Given that the inhibitory effect of memantine on AMPAR-EPSCs in VGluT2 neurons is abolished by intracellular dialysis of IEM-1460, these results support the conclusion that memantine effectively blocks postsynaptic CP-AMPARs in spinal dorsal horn neurons caused by nerve injury.

## Discussion

AMPARs are dynamic complexes capable of shifting between different subunit assemblies consisting of homo- or heteromeric combinations of GluA1 and GluA2 or GluA2 and GluA3 subunits (*1, 2*). In mature neurons, AMPARs are primarily calcium-impermeable, due to the presence of GluA2 subunits which undergo RNA editing at site 607 (Q/R) (*6–13*). In isolation, CP-AMPARs desensitize more quickly and exhibit reduced sensitivity to polyamine block and hence may not contribute significantly to an increase in calcium influx. However, when these receptors are in complex with auxiliary subunits such as γ2 and γ8 they have slower desensitization, higher residual currents, and higher single-channel conductance, leading to increased Ca^2+^ influx. This increase in calcium permeability contributes to neuronal excitability, synaptic plasticity, and neuronal survival. Thus, targeting CP-AMPARs is a critical therapeutic strategy.

Here we show that memantine, an FDA-approved drug classified as an uncompetitive antagonist of NMDARs, also inhibits CP-AMPARs. Through electrophysiology and cryo-EM, we demonstrate that memantine acts as an open channel blocker and shares similarities with the mechanism of NMDAR inhibition, such as its voltage dependence and steric effects (*15, 16*). However, in contrast to the memantine inhibition in NMDA receptors, inhibition of CP-AMPAR currents does not show the biexponential recovery from inhibition as well as pH effect of this recovery, thus suggesting that the second membrane pathway seen in NMDARs is not seen in AMPAR inhibition by memantine.

Through cryo-EM on CP-AMPARs activated in the presence of memantine, we precisely delineate the blocking mechanism. CI-AMPARs have arginine residues at the Q/R site, which dramatically lowers the memantine inhibition (Fig. 6A). CP-AMPARs, which have glutamine at the Q/R site, have significantly increased affinity for memantine, which directly interacts with the Q/R site (Fig. 6B). This causes rearrangement of the Q/R site, and narrows the selectivity filter below. Memantine’s hydrophic tricyclodecane cage sits in the hydrophobic cavity of the ion channel, immediately above the Q/R site. This likely prevents the coordination of calcium ions and water around the channel gate (*34*). Importantly, a genetic mutation in GluA2 (Q607E) that causes severe neurodevelopmental disorders, occurs directly at the Q/R site. We show that memantine block is significantly increased in Q607E CP-AMPARs, suggesting its potential therapeutic utility in conditions characterized by aberrant AMPA receptor activity in some genetic channel mutations (Fig. 6C).

**Figure 6.**
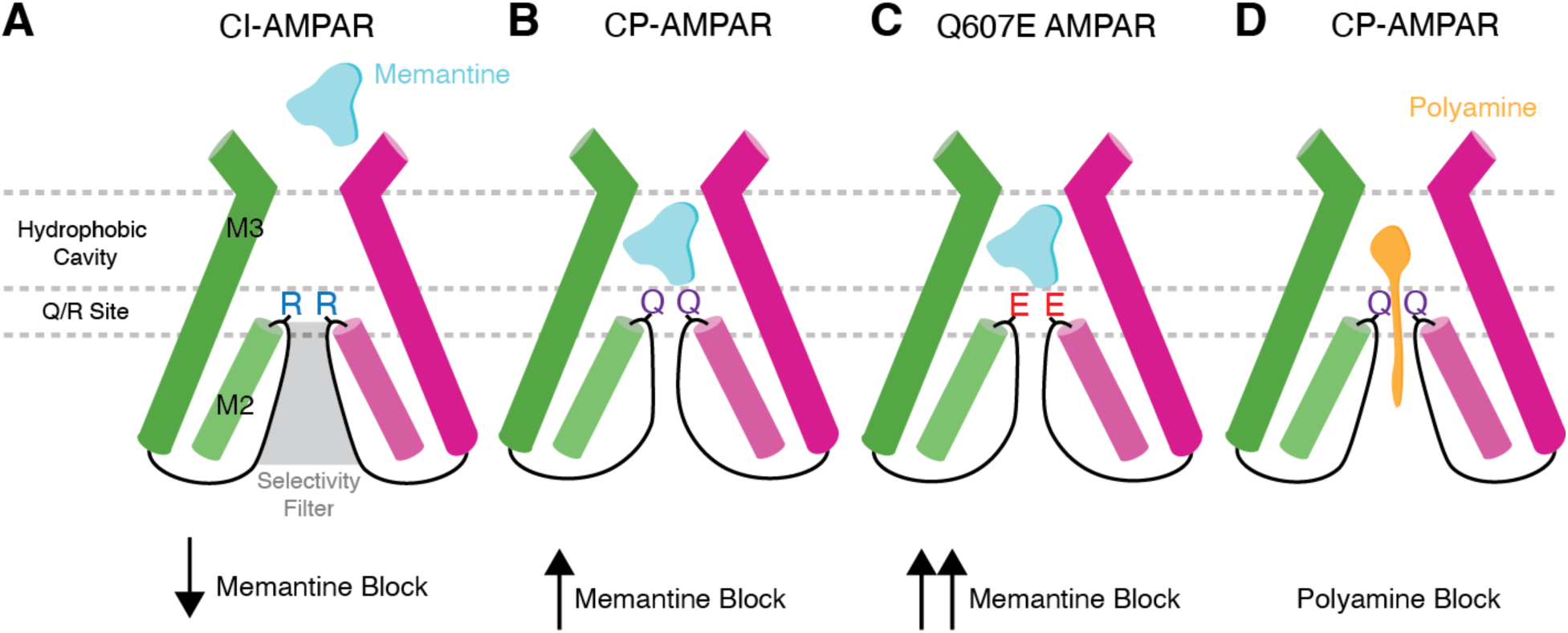
Mechanisms of channel block in AMPARs. (A) Presence of arginine at the Q/R site in CI-AMPARs dramatically reduces the efficacy of memantine bock. (B) Glutamine at the Q/R site in CP-AMPARs enables memantine inhibition in CP-AMPARs by directly coordinating memantine in the hydrophobic pocket. Memantine binding also narrows the selecitivity filter. (C) The Q607E mutation increases memantine’s inhibition efficacy. (D) Polyamines block CP-AMPARs through coordination of the polyamine tail in the selectivity filter and the hydrophobic heads being coordinated in the channel hydrophobic cavity.

Beyond its effects on isolated receptors, we also studied memantine’s impact on synaptic transmission in cultured neurons and in slice recordings from a model of nerve injury-induced neuropathic pain where synaptic CP-AMPARs are enhanced. These studies show that memantine does not have a significant effect on healthy neurons. Memantine inhibition is augmented when unedited GluA2 or GluA2 607E mutant subunits are expressed. Our experiments were done at - 60 mV and it has been shown that neurons exhibit a range of resting potentials from -60 mV to - 85 mV. We demonstrate that memantine inhibition is higher at hyperpolarized voltages. Furthermore, we show that memantine inhibits postsynaptic CP-AMPARs that are potentiated by nerve injury. Thus, we expect memantine to have inhibitory effects in pharmacologically relevant concentrations.

We also show that other NMDAR blockers such as ketamine or MK801 do not show significant inhibition of CP-AMPARs in the concentration range where they inhibit NMDARs. This adds possible pathway that may contribute to the differences between the two inhibitors that needs to be further explored.

The cryo-EM data also shows memantine block in CP-AMPARs is unique from polyamine block in CP-AMPARs. Polyamines derivates partially permeate the channel where the polyamines act as cations that are coordinated by the selecitivity filter. A hydrophobic head above the polyamines sits in the channel hydrophobic cavity, effectively plugging the channel (Fig. 6D). This is in contrast to memantine, which interacts with the Q/R site directly, narrows the selectivity filter, and blocks the channel.

In conclusion, our study presents a comprehensive examination of memantine’s pharmacological profile, revealing its novel role as an inhibitor of CP-AMPARs. These studies provide a foundation for further drug development targeting CP-AMPARs and illuminate the polypharmacology of memantine.

## Materials and Methods

### Constructs

The rat GluA2 flip (Q) construct was used to generate the GluA2(R) and GluA2(E) constructs using standard site-directed mutagenesis. The tandem constructs for GluA2 with γ2 and γ8 were generated using a Gly-Ser linker between the C-terminus of GluA2 and the N-terminus using Gibson Assembly protocol. The mutations and integrity of the constructs were confirmed by sequencing (Genewiz).

### Cell culture

The electrophysiological experiments were performed using HEK 293T cells (American tissue culture corporation). The procedure for maintenance and transfection of the HEK-293T cells has been previously described (*20, 42*). Briefly, HEK-293 cells were grown to 40%-50% confluency in DMEM (GenDEPOT) supplemented with 10% FBS (GenDEPOT) and penicillin/streptomycin (Invitrogen) and transfected with GluA2(Q), GluA2(Q)/ γ2, GluA2(Q)/ γ8, GluA2(R), or GluA2(E) along with GFP using lipofectamine 2000 (Invitrogen). Cells were re-plated after 4-6 h at a low density in fresh media containing NBQX. For single-channel recordings, cells were grown on poly-D-lysine–coated dishes, while no poly-lysine was used for whole-cell recordings. The electrophysiological experiments were performed 24-48 h after transfection.

For primary neuronal cells, the hippocampus of E-18 prenatal (Sprague-Dawley rats) embryos was dissected and dissociated as previously described (Carrillo et al., 2020a). The neurons were grown in poly-L-lysine– and laminin (Sigma-Aldrich)–coated glass coverslips, in Neurobasal medium with B27 at a high density. Fifty percent of the culture media was exchanged every two days. Primary neuronal cells (15 DIV) were transferred to serum free media 2–4 h prior to transfection. We used a DNA/Lipofectamine 3000 reagent ratio of 1 µg/1.5 µl and a DNA/P3000 ratio of 1 µg/2 µl. The neurons were seeded into round german coverslip (12 mm) and transfected using a total of 1 µg of DNA per well (GluA2(Q)/ γ2 or GluA2(E)/ γ2). After 6 h, the media was replaced for fresh media containing NBQX, after 24-48 h the electrophysiological experiments were performed. All animal experiments were conducted following the National Institutes of Health’s Guide for the care and use of laboratory animal’s guidelines, and protocols were approved by the University of Texas Health Science Center at Houston.

### Electrophysiology

Whole-cell patch-clamp recordings with HEK-293T cells were obtained from HEK293T cells using 3–5 MΩ resistance borosilicate glass pipettes. Intracellular buffer used for the whole cell recordings contained 135 mM CsF, 33 mM CsCl, 2 mM MgCl_2_,1mM CaCl_2_, 11 mM EGTA, and 10 mM HEPES, pH 7.4, while extracellular buffer contained 150 mM NaCl, 4 mM KCl, 2 mM CaCl_2_, and 10 mM HEPES, pH 7.4. Solution exchange was achieved using a perfusion Fast-Step system (Warner Instruments) with the cells being lifted and brought near the perfusion system. All recordings were performed at room temperature with a holding potential of -60 mV using an Axopatch 200B amplifier (Molecular Devices). The currents were acquired at 10 kHz using pCLAMP10 software (Molecular Devices), and filtered at 5 kHz.

For single-channel recording, the electrophysiological recordings were performed in the outside-out patch-clamp configuration, at a holding potential was -100 mV. Buffers and solution concentrations were the same as those used for whole-cell recordings. Data were acquired at 50 kHz and low-pass filtered at 10 kHz (Axon 200B and Digidata 1550A; Molecular Devices) and further filtered at 1 kHz during analysis. The recordings were idealized using the segmental k-means algorithm of QuB (*43*).

For neuronal recordings the intracellular buffer contained 120 mM Cs-gluconate, 20 mM HEPES, 4 mM MgCl_2_, 10 mM EGTA, 0.4 mM GTP-Na, 4 mM ATP-Mg, and 5 mM phosphocreatine, adjusted to pH 7.3, and the extracellular buffer contained 140 mM NaCl, 2.5 mM KCl, 2.5 mM CaCl_2_, 0.5 mM MgCl_2_, 1.25 mM NaH_2_PO_4_, 10 mM HEPES, 25 mM glucose, 1 µM Tetrodotoxin (TTX), 10 mM bicuculline, 100 µM DL-2-amino-5-phosphonopentanoic acid (DL-APV), 40 µM 5,7-dichlorokynurenic acid (DCKA) and 1 µM strychnine, adjusted to pH 7.4. Spontaneous miniature excitatory postsynaptic currents (mEPSCs) were recorded from neurons that were in culture for 14-21 d, at room temperature, using 8 – 15 MΩ resistance fire-polished borosilicate glass pipettes, with a holding potential of -80 mV using an Axopatch 200B amplifier (Molecular Devices). The currents were acquired at 50 kHz using pCLAMP10 software (Molecular Devices), and filtered online at 5 kHz. mEPSCs were analyzed using a mEPSC current-template search through Clampfit 10 software (Molecular Devices), with a detection threshold of -6 pA.

### Data analysis

Dose-response curves were generated using E_inhibition_ as a function of memantine concentrations. For these E_inhibition_ was determined using the following equation:

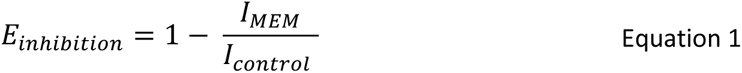

Where I_MEM_ is the steady-state current in the presence of Memantine and I_control_ is the steady-state current in the absence of Memantine.

IC_50_ was determined from the dose-response curves using the Hill equation:

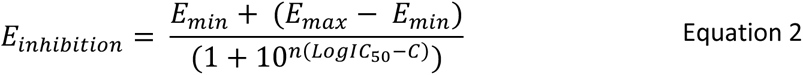

Where E_inhibition_ is steady-state inhibition as defined by Equation 1, E_min_ is the lowest value for steady-state inhibition, E_max_ is the maximum value for steady-state inhibition, IC_50_ is the concentration of Memantine at half-maximal inhibition, and C is the concentration of Memantine.

### Statistics

At least three recordings were obtained for each condition studied from at least 3 different days. All electrophysiological data were statistically analyzed using the Students’ paired t-test. These tests were performed using SigmaPlot10 (v10; Systat Software, Inc, USA). For all tests, a p-value of 0.05 was considered significant, and a p-value of 0.001 was considered highly significant.

### Sources of Drugs

Memantine (Sigma-Aldrich), D-APV (Abcam), bicuculline (Sigma-Aldrich), DCKA (Abcam), glutamate (Sigma-Aldrich), TTX (Tocris), strychnine (Sigma-Aldrich), MK-801 (Sigma-Aldrich) and cyclothiazide (CTZ) (Sigma-Aldrich) were commercially available. *N*,*N*,*N*,3,5-pentamethyladamantan-1-ammonium iodide (trimethylmemantine, TMM) was synthesized and purified as previously reported (*27*).

### Animal models

All experimental procedure and protocols were approved by the Institutional Animal Care and Use Committee at the University of Texas MD Anderson Cancer Center and adhered to the National Institutes of Health Guide for the Care and Use of Laboratory Animals. *VGluT2-ires-Cre* knock-in mice (#028863) and *tdTomato-floxed* mice (#007909) with C57BL/6 genetic background were purchased from The Jackson Laboratory. The *VGluT2^Cre/+^:tdTomato^flox/flox^*mice were generated by crossing male *VGluT2-ires-Cre* with female *tdTomato-floxed* mice (*44, 45*). Mouse genotypes were confirmed through genotyping using ear biopsies. The specificity of tdTomato-labeled VGluT2 neurons in the spinal dorsal horn has been previously validated (*44, 46, 47*). Spared nerve injury (SNI) surgery was conducted as previously outlined (*45*). Briefly, mice were anesthetized with 2%–3% isoflurane, and an incision was made on the left lateral thigh to expose the sciatic nerve. The tibial and common peroneal nerve branches were ligated with a 6-0 silk suture and sectioned distal to the ligation sites under a surgical microscope, while leaving the sural nerve intact. Male and female mice (10–12 weeks old) were used for final electrophysiological recordings three weeks after SNI surgery and housed in groups of no more than five per cage, with ad libitum access to food and water. The animal housing facility was maintained at 24°C under a 12-hour light-dark cycle.

### Electrophysiological recordings in spinal cord slices

Mice were deeply anesthetized with 3% isoflurane, and the lumbar spinal cords were quickly promptly excised via laminectomy. Transverse slices (400 μm thick) of spinal cords were then prepared using a vibratome and submerged in sucrose-modified artificial cerebrospinal fluid saturated with 95% O_2_ and 5% CO_2_. The composition of the artificial cerebrospinal fluid was as follows (in mM): 234 sucrose, 26 NaHCO_3_, 3.6 KCl, 2.5 CaCl_2_, 1.2 MgCl_2_, 1.2 NaH_2_PO_4_, and 25 glucose. Subsequently, the slices were transferred to Krebs solution containing (in mM) 117 NaCl, 25 NaHCO_3_, 3.6 KCl, 2.5 CaCl_2_, 1.2 MgCl_2_, 1.2 NaH_2_PO_4_, and 11 glucose. All slices were incubated in a continuously oxygenated chamber for a minimum of 1 hour at 34°C before being utilized for recordings.

The spinal cord slices were carefully transferred into a recording chamber and perfused continuously with oxygenated Krebs solution at a rate of 3 ml/min at 34°C. tdTomato-tagged neurons located in lamina II were identified using an upright microscope equipped with epifluorescence and differential interference contrast optics (#BX51 WI, Olympus Optical Co.). Glass recording electrodes (with resistance ranging from 5 to 8 MΩ) were filled with an internal solution containing (in mM) 135 potassium gluconate, 5 KCl, 2 MgCl_2_, 0.5 CaCl_2_, 5 ATP-Mg, 0.5 Na_2_-GTP, 5 EGTA, 5 HEPES, and 10 lidocaine N-ethyl bromide (7.3 pH, 280‒300 mOsm). In certain slice recordings, 10 mM IEM-1460 was incorporated into the intracellular solution to inhibit postsynaptic CP-AMPARs, as we described previously (*11*). Excitatory synaptic currents (EPSCs) were recorded at a holding potential of –60 mV. AP5 (50 µM) was applied to the bath solution throughout the recording period to block NMDARs. To induce the release of glutamate from primary afferents, EPSCs of labeled neurons were evoked by electrical stimulation (0.6 mA, 0.5 ms, and 0.1 Hz) of the ipsilateral dorsal root using a bipolar tungsten electrode. Monosynaptic EPSCs were identified by their consistent latency and the lack of conduction failure during 20-Hz stimulation (*44, 45*). Signal filtering was set at 1–2 kHz, and all signals were processed through a Multiclamp 700B amplifier (Molecular Devices) before being digitized at 20 kHz using DigiData 1550B (Molecular Devices). The peak amplitude of evoked EPSCs was quantified by using pClamp software (Molecular Devices). AP5 (#HB0252) was purchased from Hello Bio, and IEM-1460 (#15623) was acquired from Cayman Chemical.

### Protein expression and purification

GluA2-γ2_EM_ bacmid and P1 baculovirus were prepared as previously described (*28, 30–33*). Protein expression in mammalian Expi293F GNTI^-^ cells (Gibco, A39240) was induced by the addition of P1 baculovirus in a 1:10 ratio of P1 virus to culture volume. Cells were grown at 37°C in 5% CO_2_. 10 mM sodium butyrate (Sigma, 303410) and 2 µM ZK 20075 (Tocris, 2345) were added to cells 12 – 24 hours post-induction, then transferred to 30°C in 5% CO_2_. Cells were harvested 72 hours post-induction by centrifugation (5,000*g*, 20 minutes at 4°C), and washed with 1X PBS (pH 7.4) with protease inhibitors (0.8 µM aprotinin, 2 µg ml^-1^ leupeptin, 2 µM pepstatin A, and 1 mM phenylmethylsulfonyl fluoride). Pellets were stored at -80°C until purification. Pellets were thawed at 22°C, rotating in lysis buffer (150 mM NaCl, 20 mM Tris pH 8.0) with protease inhibitors, described above. Cells were lysed with a blunt probe sonicator (1s on, 1s off for 1 minute, 20W power, total of 3 cycles). Lysed cells were clarified by centrifugation (4,800*g*, 20 minutes at 4°C). Supernatant was collected and ultracentrifugized to isolate membranes (125,000*g*, 50 minutes at 4°C). Membrane fraction was first homogenized (Fisherbrand 150 Handheld Homogenizer) with 150 mM NaCl and 20 mM Tris pH 8.0, and then solubilized with 150 mM NaCl, 20 mM Tris pH 8.0, 1% *n*-dodecyl-ß-D-maltopyranoside (DDM; Anatrace, D310) and 0.2% cholesteryl hemisuccinate Tris salt (CHS; Anatrace, CH210), via constant stirring for 2 hours at 4°C. Sample was ultracentrifugized (125,000*g*, 50 minutes at 4°C) to separate solubilized protein. Supernatant was incubated with 1.125 ml of Strep-Tactin XT 4Flow resin (IBA, 2-5010) per 1L of cells overnight, rotating at 4°C. The resin was washed with 10 column volumes of 150 mM NaCl, 20 mM Tris pH 8.0, and 0.01% glycol-disgenin (GDN; Anatrace, GDN101) buffer. Sample was eluted with the same buffer, supplemented with 50 mM biotin. Eluate was concentrated to 500 µl volume. eGFP and Strep Tag II were then cleaved by addition of thrombin (1:200 w/w), and incubated at 22°C for 1 hour. The sample was then loaded onto a size-exclusion chromatography column (Superose 6 Increase 10/300; Cytiva, 29091596) that was equilibrated with 150 mM NaCl, 20 mM Tris pH 8.0, and 0.01% glycol-disgenin (GDN) buffer. Peak fractions were pooled and concentrated to 4 mg ml^-1^.

### Cryo-EM sample preparation and data collection

UltrAuFoil 200 mesh R 2/2 grids (Electron Microscopy Services, Q250AR2A) were plasma treated in a Pelco Easiglow (25 mA, 120 s glow time and 20 s hold time; Ted Pella, 91000). Purified sample at 4 mg ml^-1^ was supplemented with 100 µM CTZ and 500 µM memantine before ultracentrifugation (75,000*g*, 45 minutes at 4°C). Immediately prior to plunge-freezing, the sample was spiked with 1 mM Glu (pH 7.4). 3 µL of reaction mixture was applied to each grid. Grids were prepared using an FEI Vitrobot Mark IV (Thermo Fisher Scientific; wait time, 25 s; blot force, 8; blot time, 4 s) at 8°C and 100% humidity. Grids were imaged with a 300-kV Titan Krios 3i microscope equipped with a Falcon 4i camera and a Selectric energy filter set to 10-eV slit width. Micrographs were collected with a dose rate of 8.64 e^-^ /pixel/s and a total dose of 40.00 e^-^ /Å^2^. We collected a total of 7,009 micrographs (0.93 Å/pixel). Automated collection was performed with EPU software from Thermo Fisher Scientific.

### Cryo-EM Image Processing & Model Building

Cryosparc was used for all aspects of cryo-EM image processing. Refer to figure S4 and Table 1 for details. All aspects of modeling, refinement, and analysis were performed with ChimeraX, Isolde, Coot, and Phenix made accessible through the SBgrid consortium (*48–53*). See Table 1 for details. The GluA2-γ2_EM_ activated state (pdb 5WEO) was used as a starting model. Model quality was assessed with MolProbity (*54*). Pore measurements were made with HOLE (*55*).

## Acknowledgments

This study was supported by National Institute of Health Grants R35 GM122528 to V.J, NIH F99NS130928 to C.U.G, NS101880 to H-L. P. E.C.T is supported by the Searle Scholars Program (Kinship Foundation #22098168) and the Diana Helis Henry Medical Research Foundation (#142548). Cryo-EM data was collected at the Beckman Center for Cryo-EM at Johns Hopkins.

## Author contributions

Elisa Carrillo (memantine electrophysiology study design, electrophysiological recordings, data analysis, and writing). Alejandra Montaño Romero (cryo-EM study design, protein purification, cryo-EM, data analysis, writing). Cuauhtemoc U. Gonzalez (cloning and cell culture). Andreea L. Turcu (design and synthesis of Trimethylmemantine). Shao-Rui Chen (surgery, slice recording, data analysis). Hong Chen (mouse breeding and slice recording).Yuying Huang (slice recording). Hui-Lin Pan (study design of nerve injury model and writing). Santiago Vázquez (design of Trimethylmemantine and writing). Edward C. Twomey (cryo-EM study design, data analysis, writing). Vasanthi Jayaraman (memantine electrophysiology study design and writing).

## Data Availability

Cryo-EM maps and structural coordinates will be deposited into the electron microscopy data bank (EMDB) and protein data bank (pdb), respectively, upon publication.

## Supplementary Information

**Fig. S1.**
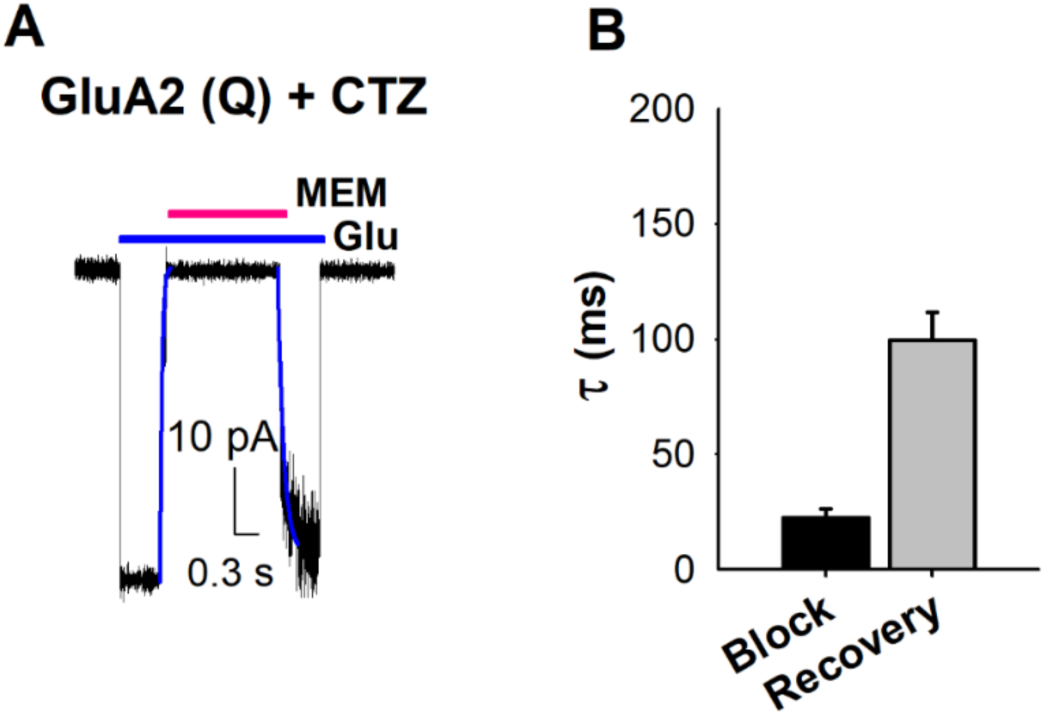
Time course of memantine inhibition. (A) Time course of the inhibition and recovery by 500 µM of memantine (MEM) in the presence of 10 mM glutamate and 100 μM CTZ. The inhibition and recovery phases were fitted to a single exponential function. (B) Bar graph showing the fits for the inhibition and recovery of memantine inhibition (n=6).

**Fig. S2.**
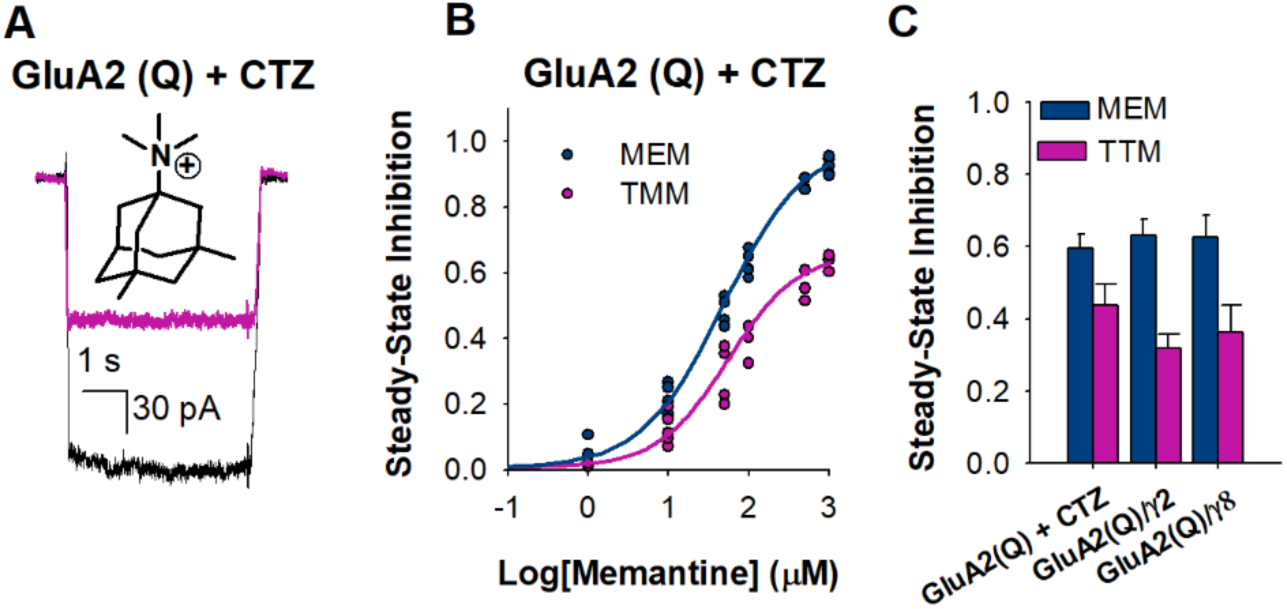
Memantine versus TMM inhibition of CP-AMPARs. (A) Chemical structure of trimethylmemantine (TMM), and representative GluA2 (Q) + CTZ current traces due to 10 mM glutamate in the absence (black) and presence of 500 µM TMM (pink). (B) The dose-dependent inhibitory effects of memantine (MEM) (•) and TMM (•) on GluA2 (Q) in the presence of CTZ, with IC_50_ 48 ± 3 µM and 384 ± 8 µM, respectively. Each dot represents data from a different cell. (C) Comparison of inhibition by 100 µM of Memantine and TMM inhibition for GluA2(Q)+CTZ, GluA2(Q)/γ2, and GluA2(Q)/γ8, (n ≥ 4).

**Figure S3.**
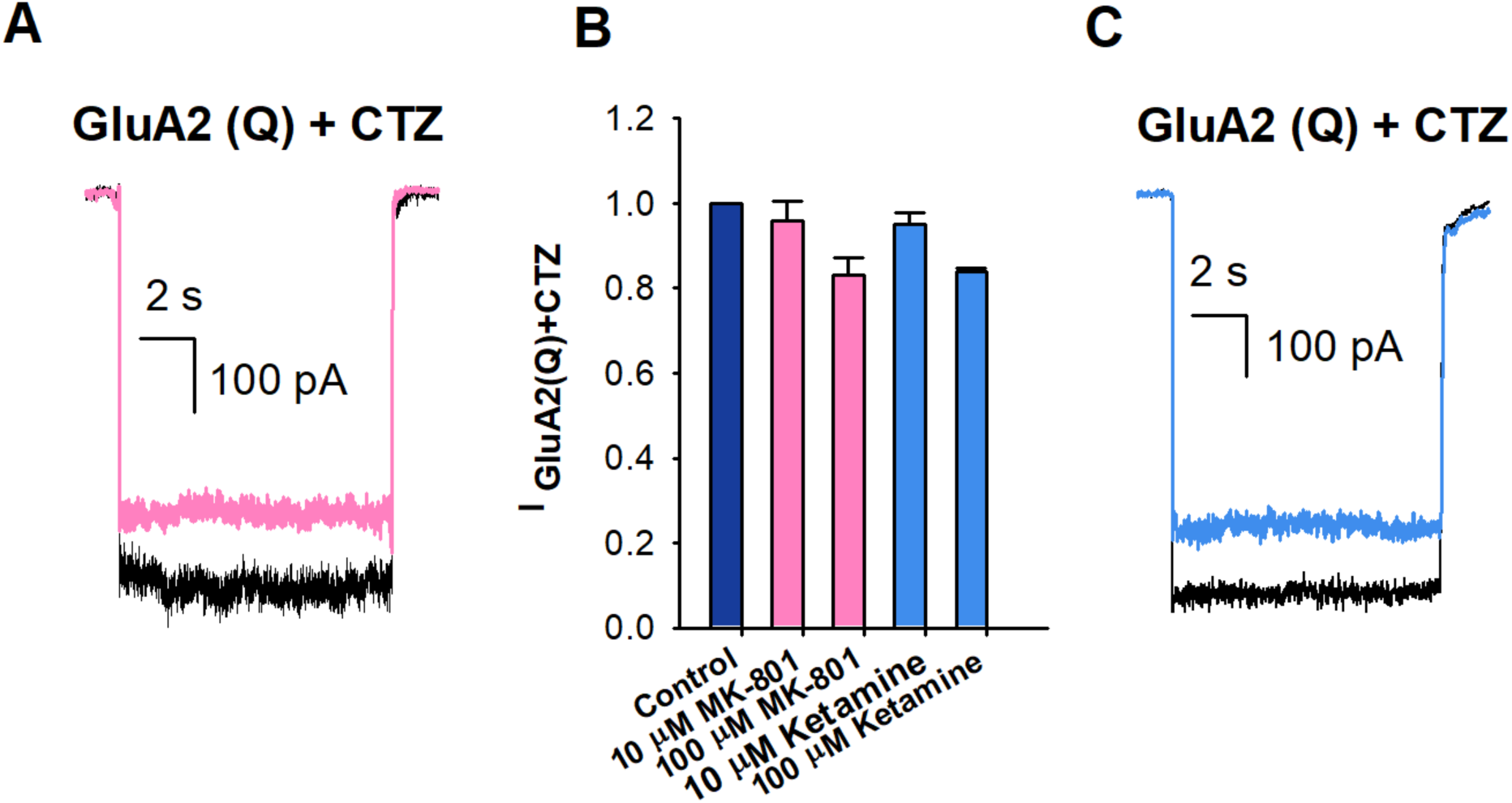
MK-801 and Ketamine responses to GluA2 (Q). (A) Representative whole-cell recordings in response to 10 mM glutamate alone (black) or in the presence of 100µM of MK-801 (pink). (B) Comparison of inhibition of 10 mM glutamate (dark blue) by 10 and 100 µM of MK-801 (pink); and 10 and 100 µM of Ketamine (blue). (C) Representative whole-cell recordings in response to 10 mM glutamate alone (black) or in the presence of 100µM of Ketamine (blue). (n ≥ 4)

**Fig. S4.**
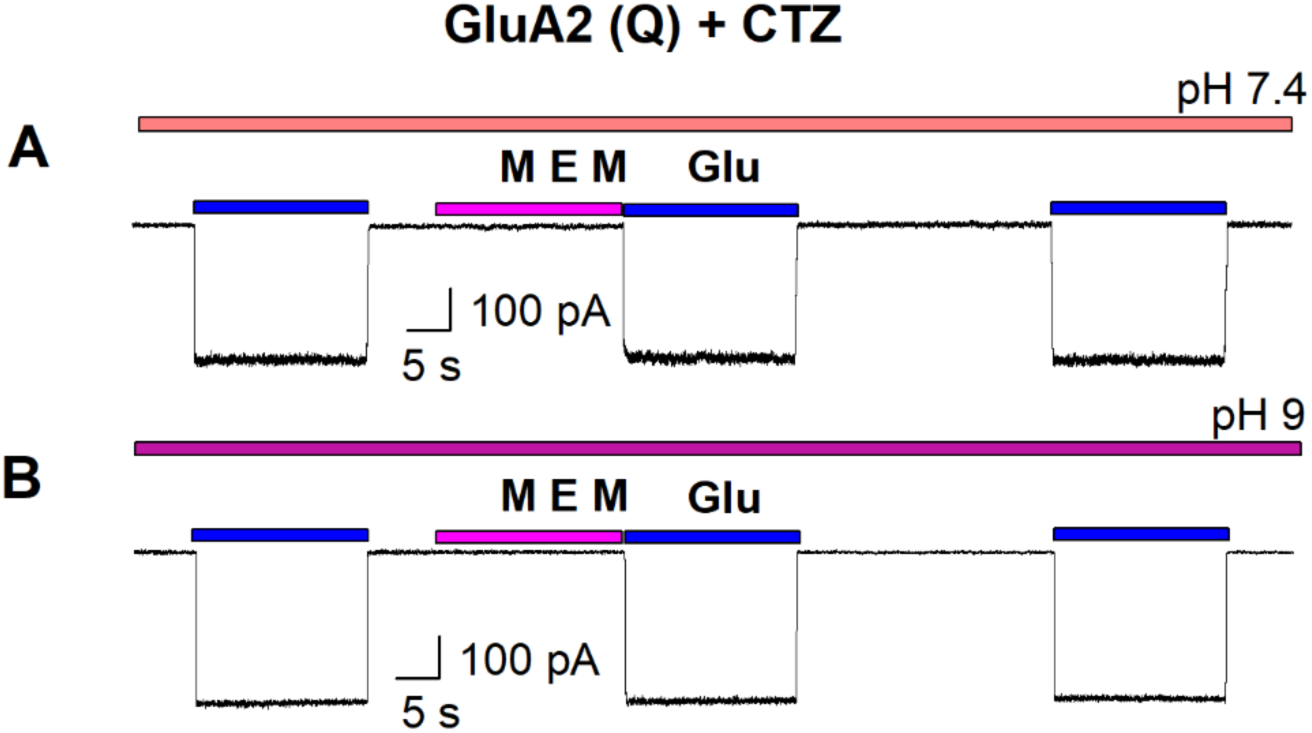
Representative current traces of memantine (MEM) inhibition showing rapid on and off rates. Current traces of memantine inhibition showing rapid on and off rates at pH 7.4 (A) and at pH 9.0 (B).

**Fig. S5.**
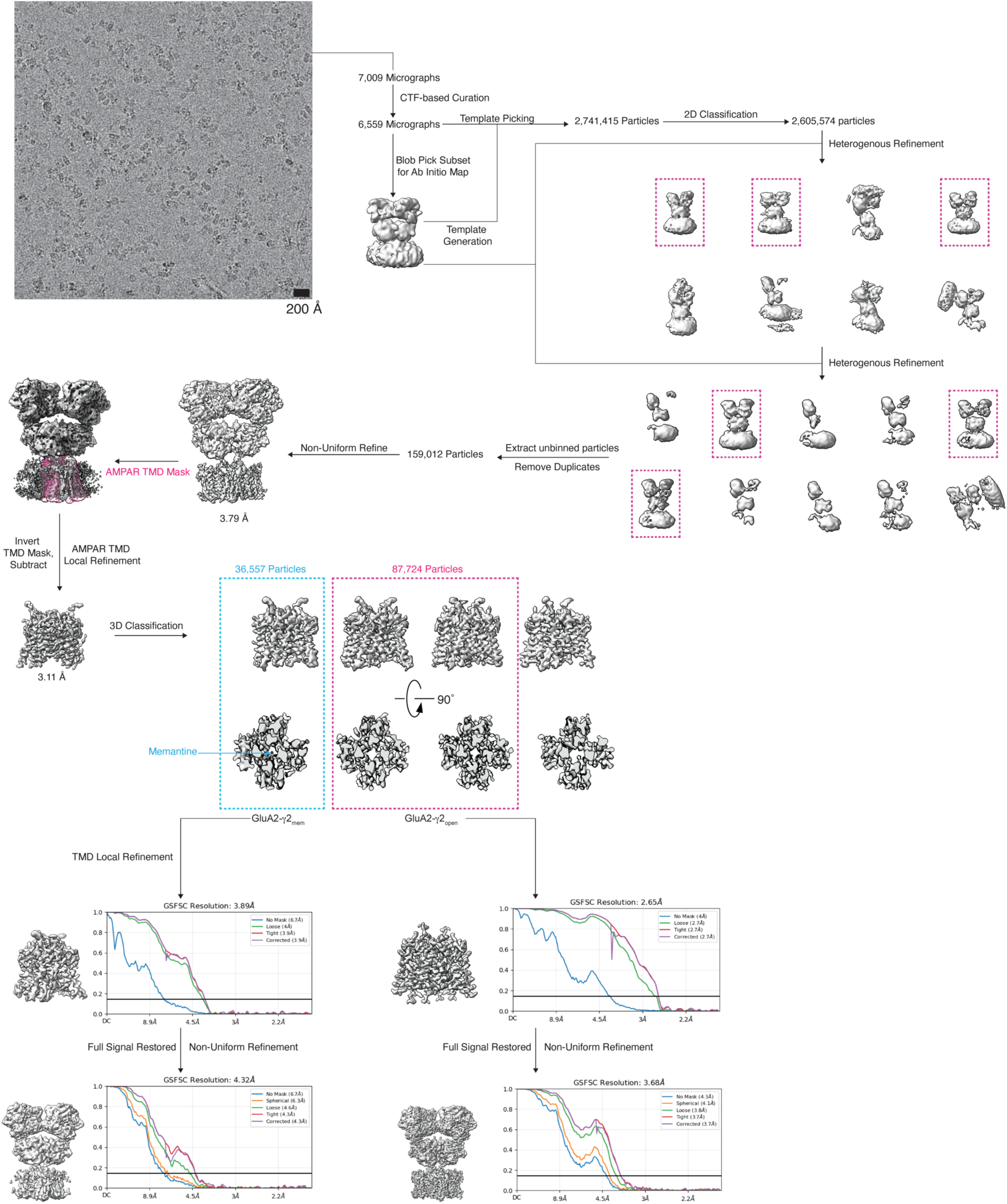
Cryo-EM processing workflow in Cryosparc.

**Figure S6.**
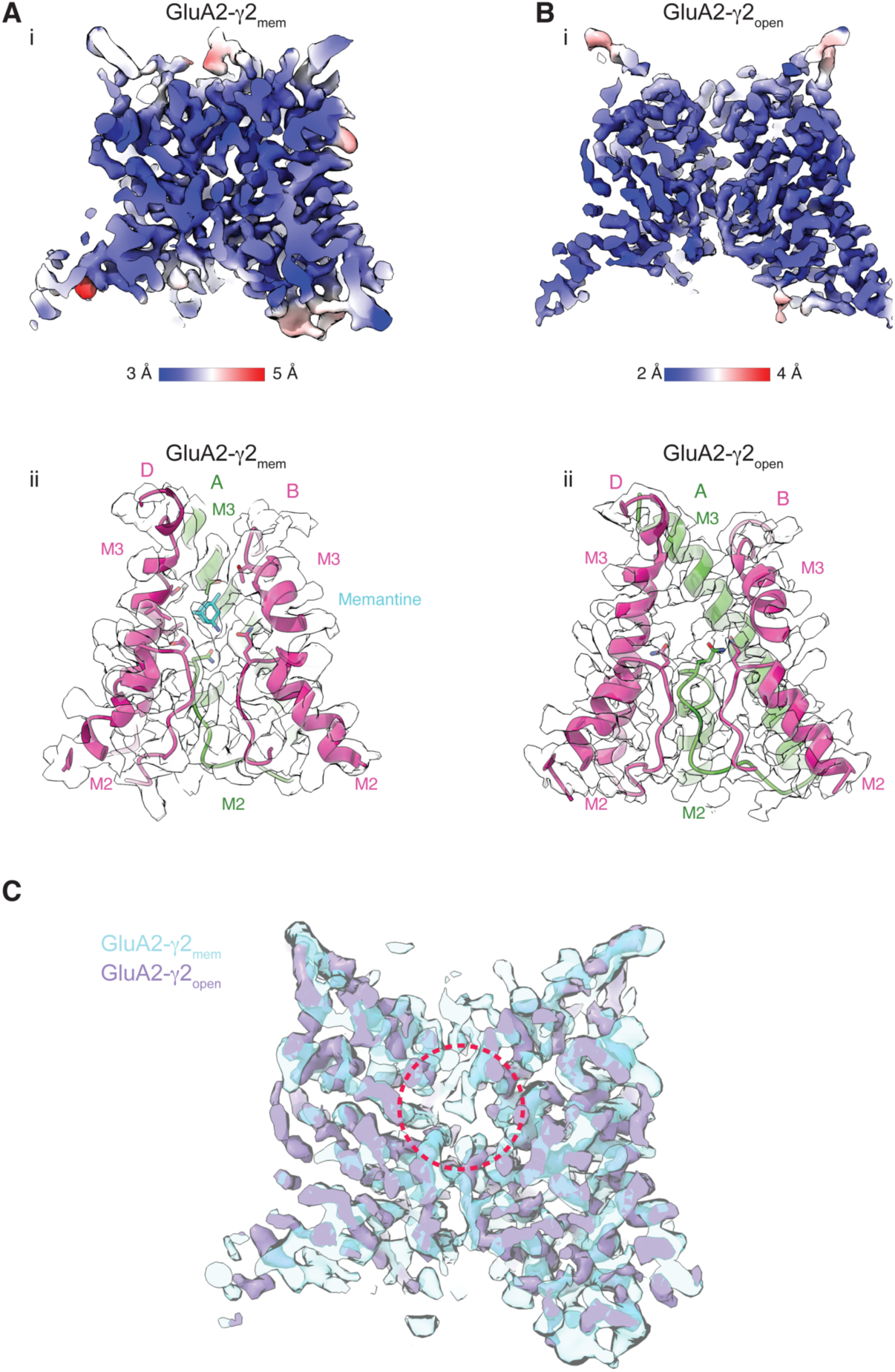
Local refinement maps and details for GluA2-γ2_mem_ and GluA2-γ2_open_. (A) Inset i, GluA2-γ2_mem_ local resolution map, colored blue (3.0 Å) to red (5.0 Å). Inset ii, GluA2-γ2_mem_ pore model fit into locally refined map. Subunit C omitted for clarity. (B) Inset i, GluA2-γ2_open_ local resolution map, colored blue (2.0 Å) to red (4.0 Å). Inset ii, GluA2-γ2_open_ pore model fit into locally refined map. Subunit C is omitted for clarity. (C) Overlay of GluA2-γ2_mem_ (cyan, transparent) and GluA2-γ2_open_ (purple) local maps. Red dashed circle indicates the memantine binding site.

**Figure S7.**
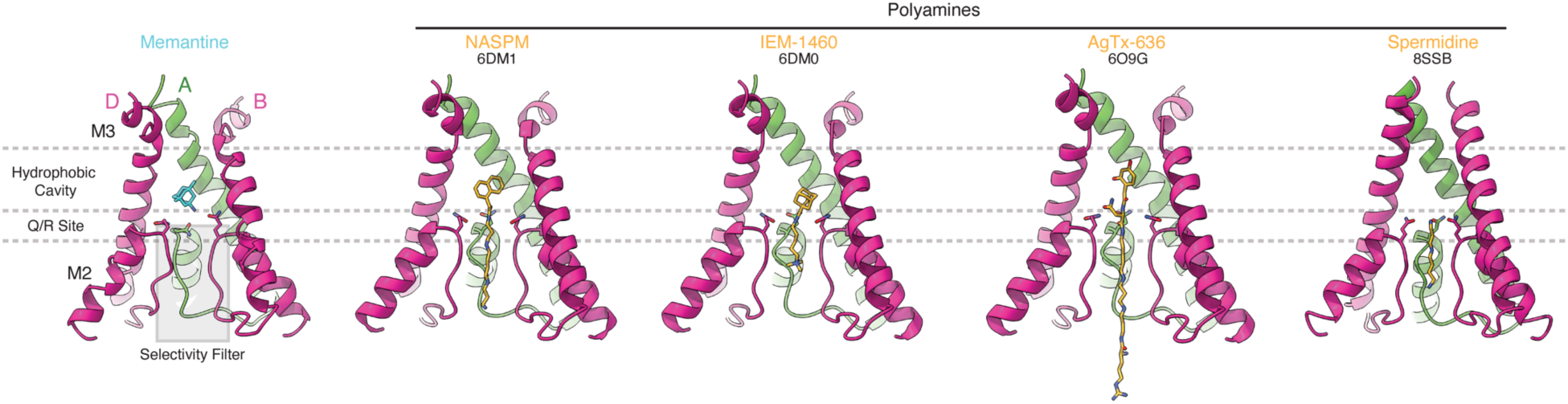
Comparison of memantine block to polyamine block. Polyamines are shown in yellow, memantine in cyan. Both are shown as sticks. Carbon molecules are colored the same as the molecule, nitrogen atoms blue, oxygen atoms red. Nitrogens in polyamine tails are directly coordinated by the selectivity filter and Q/R site, and polyamine derivates or toxins (e.g., NASPM – pdb 6DM1, IEM-1460 – pdb 6DM0, AgTx-636 – pdb 6O9G) have a hydrophobic head above the polyamine tail that sits in the hydrophobic cavity. Spermidine (pdb 8SSB) sits directly at the Q/R site and within the selecitivty filter below.

**Fig. S8.**
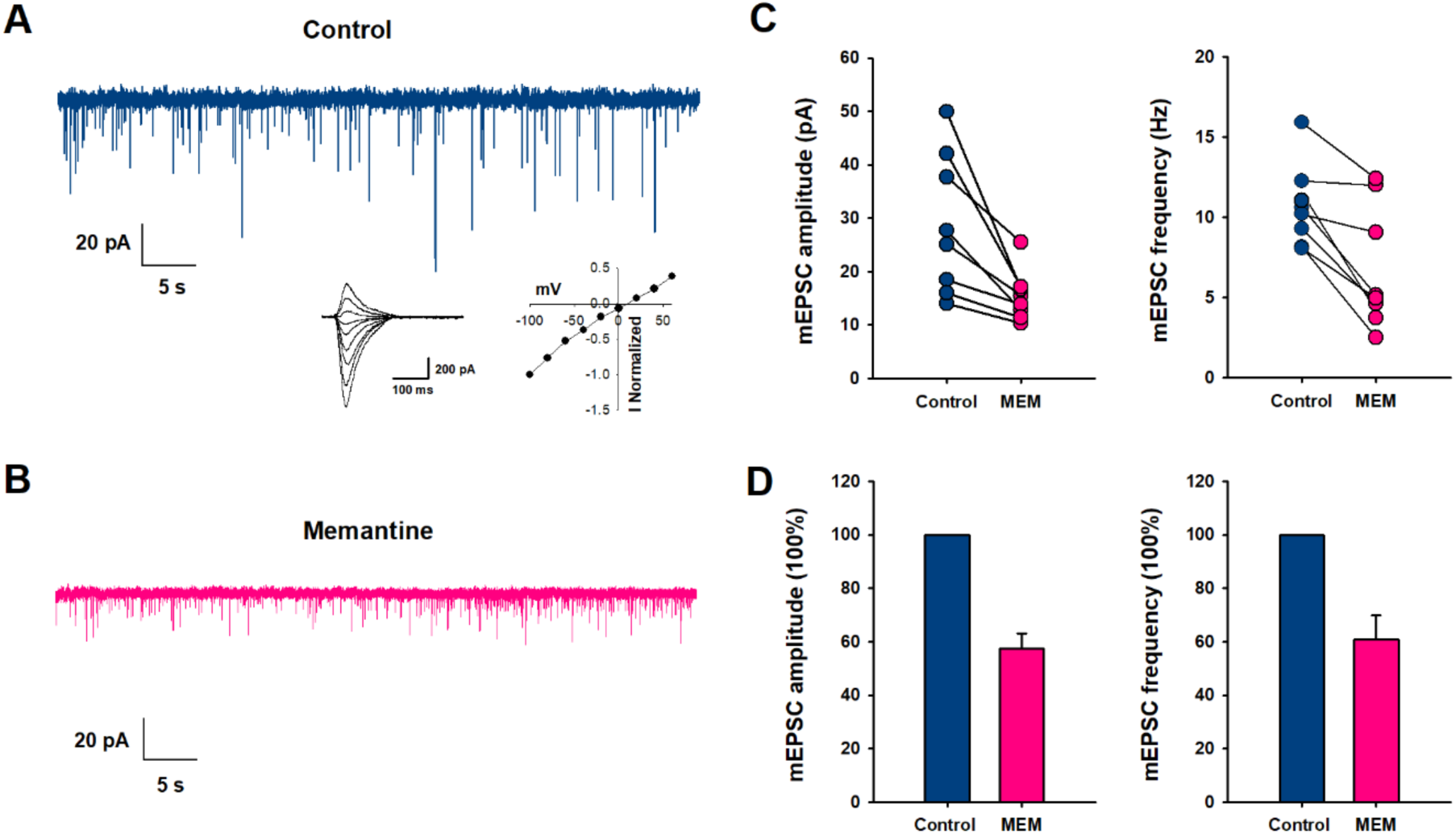
mEPSCs inhibition by 500 µM of Memantine (MEM). (A) Representative spontaneous mEPSCs, in control (blue) and (B) in the presence of 500 µM of Memantine (pink). Inset: Representative currents activated by fast application of 10 mM of glutamate (from -100 to +60 mV) from hippocampus neurons. (C) mEPSC amplitude and frequency were measured from individual neurons. Paired data from each experiment are connected by a line. (D) Bar graphs of the average values of the normalized mEPSC amplitude and frequency, in control (blue) and in the presence of 500 µM of Memantine (pink).

**Figure S9.**
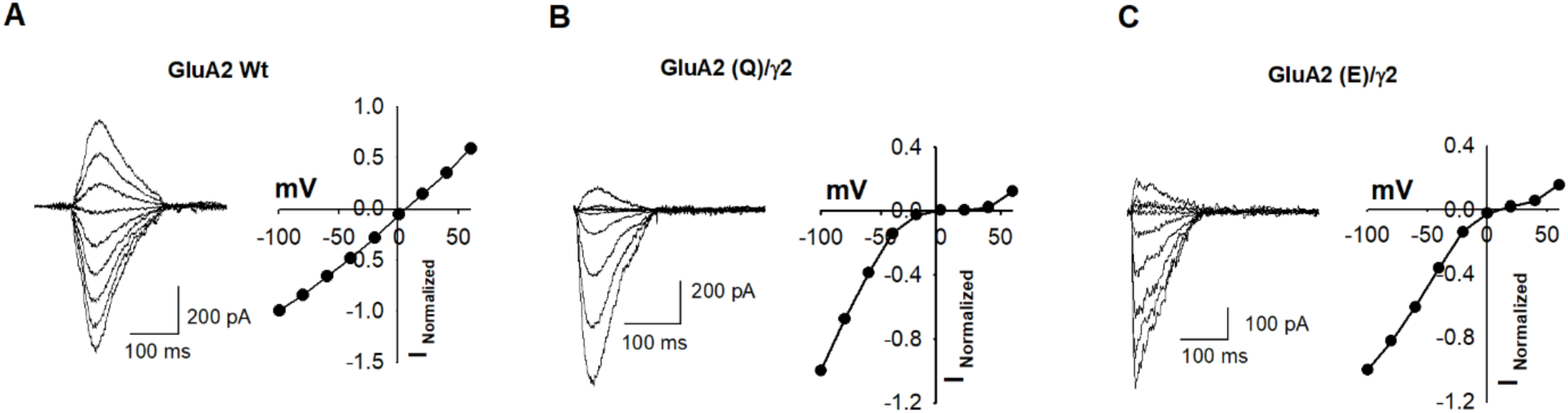
Rectification of synaptic AMPA receptors. Representative currents activated by fast application of 10 mM of glutamate (from -100 to +60 mV) from hippocampus neurons in native conditions (A), and hippocampus neurons transfected with GluA2 (Q)/γ2 (B) and GluA2 (E)/γ2 (C).

**Table S1.**
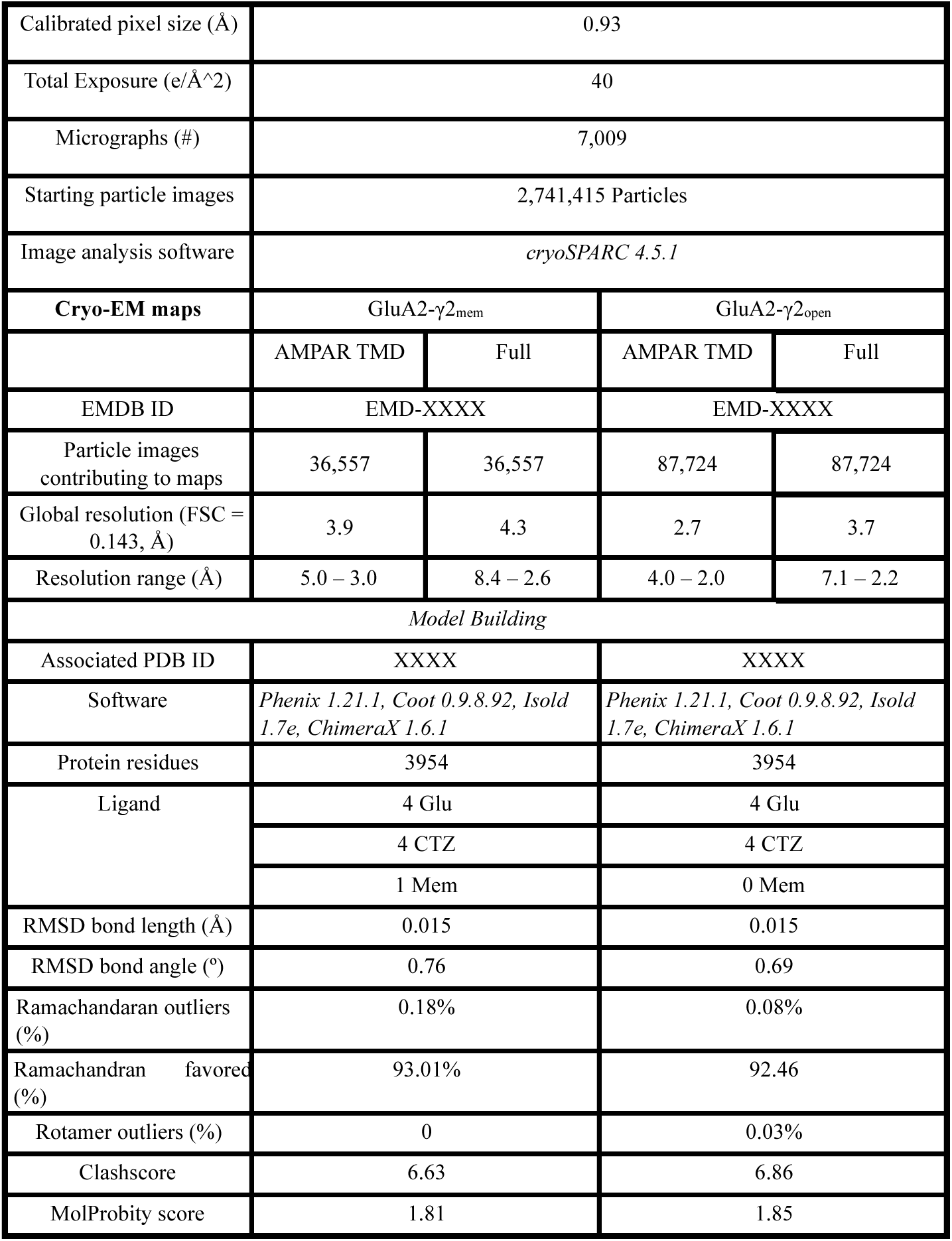

|  |  |  |  |  |
| --- | --- | --- | --- | --- |
| Calibrated pixel size (Å) | 0.93 |  |  |  |
| Total Exposure (e/Å<sup>2</sup>) | 40 |  |  |  |
| Micrographs (#) | 7,009 |  |  |  |
| Starting particle images | 2,741,415 Particles |  |  |  |
| Image analysis software | *cryoSPARC 4.5.1* |  |  |  |
| **Cryo-EM maps** | GluA2-γ<sub>2</sub><sub>mem</sub> |  | GluA2-γ<sub>2</sub><sub>open</sub> |  |
|  | AMPAR TMD | Full | AMPAR TMD | Full |
| EMDB ID | EMD-XXXX |  | EMD-XXXX |  |
| Particle images contributing to maps | 36,557 | 36,557 | 87,724 | 87,724 |
| Global resolution (FSC = 0.143, Å) | 3.9 | 4.3 | 2.7 | 3.7 |
| Resolution range (Å) | 5.0 – 3.0 | 8.4 – 2.6 | 4.0 – 2.0 | 7.1 – 2.2 |
| *Model Building* |  |  |  |  |
| Associated PDB ID | XXXX |  | XXXX |  |
| Software | *Phenix 1.21.1, Coot 0.9.8.92, Isold 1.7e, ChimeraX 1.6.1* |  | *Phenix 1.21.1, Coot 0.9.8.92, Isold 1.7e, ChimeraX 1.6.1* |  |
| Protein residues | 3954 |  | 3954 |  |
| Ligand | 4 Glu |  | 4 Glu |  |
|  | 4 CTZ |  | 4 CTZ |  |
|  | 1 Mem |  | 0 Mem |  |
| RMSD bond length (Å) | 0.015 |  | 0.015 |  |
| RMSD bond angle (°) | 0.76 |  | 0.69 |  |
| Ramachandran outliers (%) | 0.18% |  | 0.08% |  |
| Ramachandran favored (%) | 93.01% |  | 92.46 |  |
| Rotamer outliers (%) | 0 |  | 0.03% |  |
| Clashscore | 6.63 |  | 6.86 |  |
| MolProbity score | 1.81 |  | 1.85 |  |

## References

1. K. B. Hansen et al., Structure, Function, and Pharmacology of Glutamate Receptor Ion Channels. Pharmacol Rev 73, 298–487 (2021).

2. E. C. Twomey, M. V. Yelshanskaya, A. I. Sobolevsky, Structural and functional insights into transmembrane AMPA receptor regulatory protein complexes. J Gen Physiol 151, 1347–1356 (2019).

3. G. J. Iacobucci, G. K. Popescu, NMDA receptors: linking physiological output to biophysical operation. Nat Rev Neurosci 18, 236–249 (2017).

4. M. Hollmann, M. Hartley, S. Heinemann, Ca2+ permeability of KA-AMPA--gated glutamate receptor channels depends on subunit composition. Science 252, 851–853 (1991).

5. N. Burnashev, H. Monyer, P. H. Seeburg, B. Sakmann, Divalent ion permeability of AMPA receptor channels is dominated by the edited form of a single subunit. Neuron 8, 189–198 (1992).

6. I. Gaisler-Salomon et al., Hippocampus-specific deficiency in RNA editing of GluA2 in Alzheimer’s disease. Neurobiol Aging 35, 1785–1791 (2014).

7. T. Yamashita, S. Kwak, Cell death cascade and molecular therapy in ADAR2-deficient motor neurons of ALS. Neurosci Res 144, 4–13 (2019).

8. L. M. Konen et al., A new mouse line with reduced GluA2 Q/R site RNA editing exhibits loss of dendritic spines, hippocampal CA1-neuron loss, learning and memory impairments and NMDA receptor-independent seizure vulnerability. Mol Brain 13, 27 (2020).

9. S. Maas, S. Patt, M. Schrey, A. Rich, Underediting of glutamate receptor GluR-B mRNA in malignant gliomas. Proc Natl Acad Sci U S A 98, 14687–14692 (2001).

10. M. Ceprian, D. Fulton, Glial Cell AMPA Receptors in Nervous System Health, Injury and Disease. Int J Mol Sci 20, 2450 (2019).

11. L. Li et al., α2δ-1 switches the phenotype of synaptic AMPA receptors by physically disrupting heteromeric subunit assembly. Cell Rep 36, 109396 (2021).

12. E. Ghirardini et al., Mutant prion proteins increase calcium permeability of AMPA receptors, exacerbating excitotoxicity. PLoS Pathog 16, e1008654 (2020).

13. B. T. Selvaraj et al., C9ORF72 repeat expansion causes vulnerability of motor neurons to Ca(2+)-permeable AMPA receptor-mediated excitotoxicity. Nat Commun 9, 347 (2018).

14. A. L. Sladek, S. Nawy, Ocular Hypertension Drives Remodeling of AMPA Receptors in Select Populations of Retinal Ganglion Cells. Front Synaptic Neurosci 12, 30 (2020).

15. J. Bormann, Memantine is a potent blocker of N-methyl-D-aspartate (NMDA) receptor channels. Eur J Pharmacol 166, 591–592 (1989).

16. C. G. Parsons, R. Gruner, J. Rozental, J. Millar, D. Lodge, Patch clamp studies on the kinetics and selectivity of N-methyl-D-aspartate receptor antagonism by memantine (1-amino-3,5-dimethyladamantan). Neuropharmacology 32, 1337–1350 (1993).

17. L. Chen et al., Stargazin regulates synaptic targeting of AMPA receptors by two distinct mechanisms. Nature 408, 936–943 (2000).

18. Alexander C. Jackson, Roger A. Nicoll, The Expanding Social Network of Ionotropic Glutamate Receptors: TARPs and Other Transmembrane Auxiliary Subunits. Neuron 70, 178–199 (2011).

19. B. Herguedas et al., Mechanisms underlying TARP modulation of the GluA1/2-γ8 AMPA receptor. Nature Communications 13, 734 (2022).

20. E. Carrillo et al., Mechanism of modulation of AMPA receptors by TARP-gamma8. J Gen Physiol 152, e201912451 (2020).

21. I. D. Coombs, D. M. MacLean, V. Jayaraman, M. Farrant, S. G. Cull-Candy, Dual Effects of TARP gamma-2 on Glutamate Efficacy Can Account for AMPA Receptor Autoinactivation. Cell Rep 20, 1123–1135 (2017).

22. D. M. MacLean, S. S. Ramaswamy, M. Du, J. R. Howe, V. Jayaraman, Stargazin promotes closure of the AMPA receptor ligand-binding domain. J Gen Physiol 144, 503–512 (2014).

23. S. A. Shaikh et al., Stargazin Modulation of AMPA Receptors. Cell Rep 17, 328–335 (2016).

24. D. Zhang et al., Modulatory mechanisms of TARP γ8-selective AMPA receptor therapeutics. Nat Commun 14, 1659 (2023).

25. V. Salpietro et al., AMPA receptor GluA2 subunit defects are a cause of neurodevelopmental disorders. Nature Communications 10, 3094 (2019).

26. W. Hu, B. P. Bean, Differential Control of Axonal and Somatic Resting Potential by Voltage-Dependent Conductances in Cortical Layer 5 Pyramidal Neurons. Neuron 97, 1315–1326.e1313 (2018).

27. M. R. Wilcox et al., Inhibition of NMDA receptors through a membrane-to-channel path. Nat Commun 13, 4114 (2022).

28. W. D. Hale et al., Allosteric competition and inhibition in AMPA receptors. Nat Struct Mol Biol, (2024).

29. M. V. Yelshanskaya, D. S. Patel, C. M. Kottke, M. G. Kurnikova, A. I. Sobolevsky, Opening of glutamate receptor channel to subconductance levels. Nature 605, 172–178 (2022).

30. E. C. Twomey, M. V. Yelshanskaya, A. A. Vassilevski, A. I. Sobolevsky, Mechanisms of Channel Block in Calcium-Permeable AMPA Receptors. Neuron 99, 956–968 e954 (2018).

31. E. C. Twomey, M. V. Yelshanskaya, R. A. Grassucci, J. Frank, A. I. Sobolevsky, Structural Bases of Desensitization in AMPA Receptor-Auxiliary Subunit Complexes. Neuron 94, 569–580 e565 (2017).

32. E. C. Twomey, M. V. Yelshanskaya, R. A. Grassucci, J. Frank, A. I. Sobolevsky, Channel opening and gating mechanism in AMPA-subtype glutamate receptors. Nature 549, 60–65 (2017).

33. E. C. Twomey, M. V. Yelshanskaya, R. A. Grassucci, J. Frank, A. I. Sobolevsky, Elucidation of AMPA receptor-stargazin complexes by cryo-electron microscopy. Science 353, 83–86 (2016).

34. T. Nakagawa, X. T. Wang, F. J. Miguez-Cabello, D. Bowie, The open gate of the AMPA receptor forms a Ca(2+) binding site critical in regulating ion transport. Nat Struct Mol Biol 31, 688–700 (2024).

35. T. H. Chou et al., Structural insights into binding of therapeutic channel blockers in NMDA receptors. Nat Struct Mol Biol 29, 507–518 (2022).

36. S. R. Chen, J. Zhang, H. Chen, H. L. Pan, Streptozotocin-Induced Diabetic Neuropathic Pain Is Associated with Potentiated Calcium-Permeable AMPA Receptor Activity in the Spinal Cord. J Pharmacol Exp Ther 371, 242–249 (2019).

37. S. R. Chen, H. Y. Zhou, H. S. Byun, H. L. Pan, Nerve injury increases GluA2-lacking AMPA receptor prevalence in spinal cords: functional significance and signaling mechanisms. J Pharmacol Exp Ther 347, 765–772 (2013).

38. S. R. Chaplan, A. B. Malmberg, T. L. Yaksh, Efficacy of spinal NMDA receptor antagonism in formalin hyperalgesia and nerve injury evoked allodynia in the rat. J Pharmacol Exp Ther 280, 829–838 (1997).

39. S. R. Chen, G. Samoriski, H. L. Pan, Antinociceptive effects of chronic administration of uncompetitive NMDA receptor antagonists in a rat model of diabetic neuropathic pain. Neuropharmacology 57, 121–126 (2009).

40. G. F. Zhang et al., α2δ-1 Upregulation in Primary Sensory Neurons Promotes NMDA Receptor-Mediated Glutamatergic Input in Resiniferatoxin-Induced Neuropathy. J Neurosci 41, 5963–5978 (2021).

41. K. Koga et al., Voltage-gated calcium channel subunit alpha(2)delta-1 in spinal dorsal horn neurons contributes to aberrant excitatory synaptic transmission and mechanical hypersensitivity after peripheral nerve injury. Front Mol Neurosci 16, 1099925 (2023).

42. E. Carrillo, N. K. Bhatia, A. M. Akimzhanov, V. Jayaraman, Activity Dependent Inhibition of AMPA Receptors by Zn(2). J Neurosci 40, 8629–8636 (2020).

43. F. Qin, Restoration of single-channel currents using the segmental k-means method based on hidden Markov modeling. Biophys J 86, 1488–1501 (2004).

44. S. R. Chen, H. Chen, D. Jin, H. L. Pan, Brief Opioid Exposure Paradoxically Augments Primary Afferent Input to Spinal Excitatory Neurons via alpha2delta-1-Dependent Presynaptic NMDA Receptors. J Neurosci 42, 9315–9329 (2022).

45. Y. Huang et al., Theta-Burst Stimulation of Primary Afferents Drives Long-Term Potentiation in the Spinal Cord and Persistent Pain via alpha2delta-1-Bound NMDA Receptors. J Neurosci 42, 513–527 (2022).

46. L. Wang et al., Regulating nociceptive transmission by VGluT2-expressing spinal dorsal horn neurons. J Neurochem 147, 526–540 (2018).

47. T. J. Browne et al., Transgenic Cross-Referencing of Inhibitory and Excitatory Interneuron Populations to Dissect Neuronal Heterogeneity in the Dorsal Horn. Front Mol Neurosci 13, 32 (2020).

48. E. F. Pettersen et al., UCSF ChimeraX: Structure visualization for researchers, educators, and developers. Protein Sci 30, 70–82 (2021).

49. T. I. Croll, ISOLDE: a physically realistic environment for model building into low-resolution electron-density maps. Acta Crystallogr D Struct Biol 74, 519–530 (2018).

50. P. Emsley, K. Cowtan, Coot: model-building tools for molecular graphics. Acta Crystallogr D Biol Crystallogr 60, 2126–2132 (2004).

51. D. Liebschner et al., Macromolecular structure determination using X-rays, neutrons and electrons: recent developments in Phenix. Acta Crystallogr D Struct Biol 75, 861–877 (2019).

52. P. V. Afonine et al., Real-space refinement in PHENIX for cryo-EM and crystallography. Acta Crystallogr D Struct Biol 74, 531–544 (2018).

53. A. Morin et al., Collaboration gets the most out of software. Elife 2, e01456 (2013).

54. C. J. Williams et al., MolProbity: More and better reference data for improved all-atom structure validation. Protein Sci 27, 293–315 (2018).

55. O. S. Smart, J. G. Neduvelil, X. Wang, B. A. Wallace, M. S. Sansom, HOLE: a program for the analysis of the pore dimensions of ion channel structural models. J Mol Graph 14, 354–360, 376 (1996).

